# Broad substrate scope C-C oxidation in cyclodipeptides catalysed by a flavin-dependent filament

**DOI:** 10.1101/2024.07.24.604905

**Authors:** Emmajay Sutherland, Christopher J. Harding, Tancrede Martin Y Du Monceau De Bergendal, Gordon J. Florence, Katrin Ackermann, Bela E. Bode, Silvia Synowksy, Ramasubramanian Sundaramoorthy, Clarissa Melo Czekster

**Author notes:** To whom correspondence should be addressed: Clarissa Melo Czekster – School of Biology, University of St Andrews, Biomedical Sciences Research Complex, St Andrews, Fife KY16 9ST, U.K., Ramasubramanian Sundaramoorthy – Laboratory of Chromatin Structure and Function, MCDB, School of Life Sciences, University of Dundee, Dundee, DD1 5EH, UK. **Present Addresses** Emmajay Sutherland – University of Washington, Department of Chemistry, Seattle, Washington 98195-1700, United States.

## Abstract

Cyclic dipeptides are produced by organisms in all domains of life. Many possess biological activities as anticancer and antimicrobial compounds. Oxidations are frequently present in these biologically active peptides. C-C bond oxidation is catalysed by tailoring enzymes, including cyclodipeptide oxidases. These flavin-dependent enzymes are underexplored due to their complex three-dimensional arrangement involving multiple copies of two distinct small subunits and unclear mechanistic features underlying substrate selection and catalysis. Here, we determined the structure and mechanism of the cyclodipeptide oxidase from the halophile *Nocardiopsis dassonvillei* (*Ndas*CDO), part of the biosynthetic pathway for nocazine natural products. We show *Ndas*CDO forms filaments in solution with a covalently bound FMN cofactor in the interface between three distinct subunits. The enzyme is promiscuous, using many cyclic dipeptides as substrates in a distributive manner. The reaction is optimal at high pH, involving the formation of a radical intermediate. Pre-steady state kinetics, a sizeable solvent kinetic isotope effect and lack of viscosity effects support that a step coupled to FMN regeneration determines the rate of the reaction. Our work dissects the complex mechanistic and structural features of this dehydrogenation reaction. This sets the stage to utilizing *Ndas*CDO as a biocatalyst and expands the FMN-dependent oxidase family to include enzyme filaments.

## Introduction

Secondary metabolites of microbial origin are a vital but largely unexplored source of compounds with untapped therapeutic potential. Across all domains of life, cyclodipeptides (CDPs) exist either independently or embedded as part of larger, more complex natural products^1^. Distinguished by their 2,5-diketopiperazine core, CDPs are considered privileged frameworks that resist proteolysis, exhibit easy gut absorption, and can permeate the blood-brain barrier.^2^ These properties have since facilitated the use of CDPs as antibacterial^3^, antifungal^4^, antiviral^5^ and anticancer^2, 6^ agents. Whilst the precise functions of these molecules remain elusive, emerging evidence suggests their involvement in quorum sensing and transkingdom signalling.^7, 8^

Presently, three documented biosynthetic routes lead to a variety of cyclodipeptides: nonribosomal peptide synthetases (NRPSs)^9^, cyclodipeptide synthases (CDPSs)^10^, and the recently discovered arginine-containing cyclodipeptide synthases (RCDPSs)^11^. Biosynthetic gene clusters for CDPSs have been predominantly identified in bacteria, whereas NRPSs and RCDPSs from eukaryotes have been described. NRPSs, with their large multi-modular architecture, are adept at utilising inactivated amino acids to generate cyclodipeptide products.^12^ In contrast, both CDPSs and RCDPSs utilize aminoacylated-tRNAs as substrates, therefore competing with protein translational processes.^13^ Moreover, biosynthetic gene clusters encoding CDPSs often harbour tailoring enzymes that can further modify the CDP scaffold, commonly through methylation, prenylation and oxidation.^14^ Specifically, oxidation can be catalysed by cytochrome P450s^15^ and cyclodipeptide oxidases^16^ (CDOs, Figure S1 depicts the genomic context of *Ndas*CDO and similar biosynthetic gene clusters according to webflags^17^)

CDOs enable the facile incorporation of a Cα-Cβ double bond into the 2,5-diketopiperazine backbone (Figure 1a). Oxidized CDPs are building blocks for the natural product pulcherrimin, which acts as an iron chelator^18^, and phenylahistin, a precursor to the anticancer drug Plinabulin^19^. Synthetic routes for the generation of specific oxidized CDPs are scarce, and yield side products which then require lengthy purification steps.

**Figure 1:**
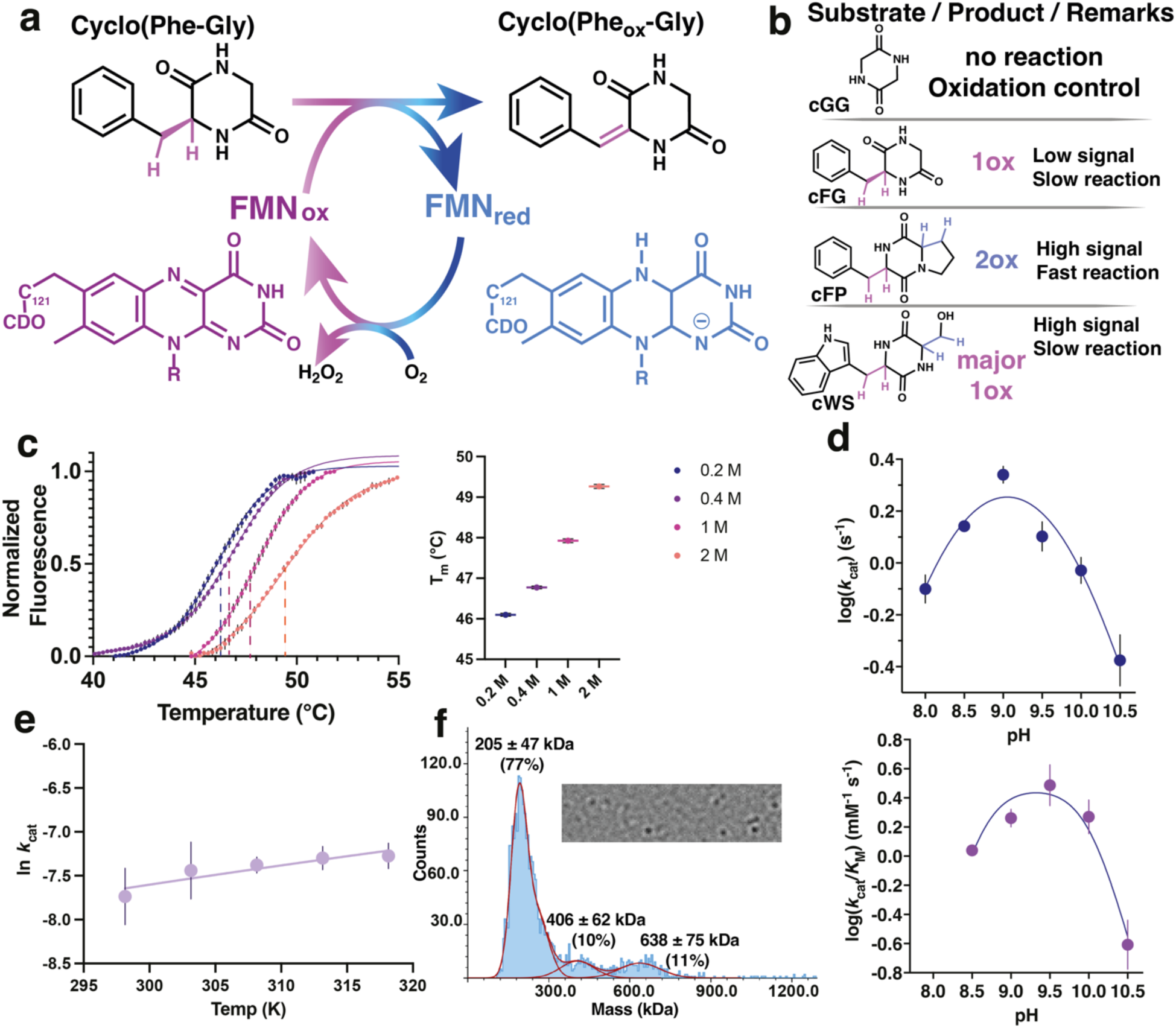
Reaction catalysed by CDO enzymes, effect of temperature, pH and oligomeric state on *Ndas*CDO. a) CDP oxidation (cFG) depicted as an example), aided by an FMN cofactor and using oxygen and a CDP as substrates to generate an oxidized CDP and hydrogen peroxide (H_2_O_2_). FMN: Flavin mononucleotide in its oxidized (FMN_ox_) or reduced (FMN_red_) forms. b) Criteria for usage of distinct CDP substrates for different experiments: cGG is a reaction control as it cannot be oxidized; catalysis with cFG is slow and a single oxidation is possible – oxidation leads to a small change in absorbance at 297 nm; cFP undergoes fast oxidation, and two oxidations are possible – oxidation leads to a large change in absorbance at 312 nm; (Hydrogen atoms depicted purple when participating in the first oxidation and blue when participating in the second oxidation rection); cWS undergoes slow oxidation, and although two oxidations are possible, second is less favourable – oxidation leads to a large change in absorbance at 297 nm; c) Melting curves using differential scanning fluorimetry (DSF), right: raw data (circles) and data fited to a Boltzmann sigmoidal equation (lines); right: fitted values for T_m_ with varying NaCl concentrations. d) pH dependence of log (*k*_cat_) and log (*k*_cat_/*K*_M_) for varied cFG (blue and purple, respectively) examined using a mixed buffer system. Data were obtained at 30 °C whilst varying the concentration of substrate. log (*k*_cat_) data are fit to a 1-proton dependence (Equation 2), log (*k*_cat_/*K*_M_) are fit to a 2-proton dependence equation (Equation 3), shown as mean values ± SEM across three replicates. Errors for *k*_cat_/*K*_M_ and *k*_cat_/*K*_ACT_ were propagated appropriately. e) Temperature dependence of *k*_cat_, data were fitted to Equation 1 yielding Δ*H*^‡^ = 4.06 ± 2.42 kcal mol^−1^ and Δ*S*^‡^ of –0.049 ± 0.008 kcal mol^−1^. f) Top, snapshot of mass photometry movie, wich was recorded for 120 s. Data was using the DiscoverMP software to generate the mass distribution histograms displayed in the bottom. Under these conditions a predominant species of 205 ± 47 kDa is observed.

Since their discovery in 2001^20^, CDOs have remained elusive, lacking structural or detailed mechanistic characterization. There are composed of two subunits – A and B – where subunit A (22 kDa) is approximately twice the size of subunit B. These, together with a covalently bound flavin cofactor comprise the catalytically competent CDO complex.^21^ Within gene clusters containing CDOs, the DNA sequences encoding subunits A and B overlap by 20-30 nucleotides, a feature postulated to act as a natural regulatory mechanism governing the expression of both subunits.^21^ Moreover, all attempts to express these subunits individually yielded inactive forms, therefore emphasising that complex formation is key to enzyme functionality.^22^ While it is established that CDOs employ molecular oxygen for dehydrogenation, producing hydrogen peroxide as a by-product, the overarching catalytic mechanism remains largely undetermined.

Notably, CDOs feature in four documented biosynthetic pathways: albonoursin from *Streptomyces noursei*^21^; nocazine from *Nocardiopsis dassonvillei*^16^; guanitrypmycin from *Streptomyces monomycini*^23^; and purincyclamide from *Streptomyces chrestomyceticus*^24^. Biocatalysis utilizing CDOs has previously involved in vivo systems, wherein media supernatants were screened for potential oxidized cyclodipeptide products^25^. Intriguingly, efforts have successfully combined CDPS and CDOs from different organisms within a singular vector, yielding a diverse array of dehydrogenated CDPs^22^. Nevertheless, a deeper molecular understanding of CDO structure and mechanism promises to enhance their utility as biocatalysts. By exploring reaction conditions to achieve optimal enzymatic efficiency whilst maintaining CDO stability, these enzymes can be harnessed as powerful tools for facilitating the introduction of C-C double bonds into cyclodipeptides.

The substrate scope of CDOs is varied with a singular CDO capable of accepting numerous CDPs. However the reaction maintains exquisite selectivity for Cα-Cβ oxidation where FMN regeneration does not require additional cofactors. Herein, we describe an in-depth characterisation of *Ndas*CDO, the cyclodipeptide oxidase found in the nocazine biosynthetic pathway of *Nocardiopsis dassonvillei*^16^. Combining a plethora of biochemical and biophysical techniques including cryo-EM, EPR and kinetic assays in steady state and pre-steady state we determine substrate scope and rate limiting steps in the reaction, unveiling crucial features of *Ndas*CDO as an FMN-dependent filament enzyme catalysing CDP dehydrogenation.

## Methods

### Chemical Reagents and General Methods

Buffers, salts, and other common chemicals were purchased from Merck or Fisher Scientific and used without further purification. All cyclodipeptide substrates (except for cWS, which was synthesized chemically as described in the supporting information) were purchased from Bachem and used without further purification. All spectrometric experiments were performed in triplicate using a BMG Labtech POLARstar Omega plate reader. All experiments in replicate were from individual measurements, and not from repeated measurements on the same sample. Errors were propagated following standard formulas for uncertainty propagation, including for all *k*_cat_/*K*_M_ values.^26^

### Expression, purification, and mutagenesis of *Ndas*CDO

Genes encoding *Ndas*1146 (Uniprot: D7B1W6) and *Ndas*1147 (Uniprot: D7B1W7) from *Nocardiopsis dassonvillei* were cloned into a pRSFDuet-1 vector by Genscript. Before further use, the plasmid was transformed into commercially available *Escherichia coli* (*E. coli*) DH5α cells (NEB). The DNA was extracted following the QIAprep Spin Miniprep Kit (Qiagen) and sequenced by DNA Sequencing & Services (MRC I PPU, School of Life Sciences, University of Dundee, Scotland, www.dnaseq.co.uk) using Applied Biosystems Big-Dye Ver 3.1 chemistry on an Applied Biosystems model 3730 automated capillary DNA sequencer. Mutants of *Ndas*CDO were created by site-directed mutagenesis based on the NEB Q5 site-directed mutagenesis kit (Table S1 for primer details).

For protein expression, purified plasmid was transformed into *E. coli* BL21(DE3) competent cells (NEB). Cells were grown at 37 °C at 180 rpm until an optical density (OD_600_) between 0.6 and 0.8 was reached. Protein expression was induced upon the addition of IPTG to a final concentration of 500 μM. The cells were incubated overnight at 25 °C before being harvested via centrifugation. Cells were resuspended in lysis buffer consisting of 100 mM Tris pH 8, 200 mM NaCl, 20 mM imidazole, 10% (v/v) glycerol and stirred to homogeneity in the presence of lysozyme (10 mg) and DNase I (1 mg) at 4 °C. Cells were lysed at high pressure using a cell disrupter at 30 kpsi and the lysate was clarified via centrifugation. The resultant supernatant was filtered through a 0.8 μm membrane before being loaded onto 5 mL HisTrap FF nickel column (GE Healthcare) pre-equilibrated in lysis buffer. Proteins were eluted in steps of 10%, 20% and 100% elution buffer containing 100 mM Tris pH 8, 200 mM NaCl, 300 mM imidazole, and 10% (v/v) glycerol. Fractions of interest were pooled together for dialysis into 50 mM Tris pH 8, 200 mM NaCl, 10% (v/v) glycerol overnight at 4 °C. The following day, the protein was concentrated using an Amicon Stirred Cell (Merck) equipped with a 100 kDa molecule weight cut off (MWCO) ultrafiltration disc (Millipore). To concentrate to a final smaller volume, the protein was transferred to a Vivaspin® 6 centrifuge unit with a 100 kDa MWCO PES membrane (Sartorius).

### Melting temperature investigations of *Ndas*CDO

Differential scanning fluorimetry was performed in triplicate using a Quantstudio real time thermal cycler (Thermofisher). SYPRO™ Orange Protein Gel Stain (Thermofisher) was used to a final concentration of 10x with 50 μM *Ndas*CDO in 20 μL. Buffer and salt conditions were varied using a series of conditions prepared in an in-house stability screen, and the temperature was ramped from 25 to 95 °C over 90 minutes. Melting temperatures were calculated by fitting the data to the Boltzmann sigmoidal function (Graphpad Prism).

### Mass photometry

Mass photometry experiment was performed using a OneMP mass photometer (Refeyn Ltd). Microscope slides (70 x 26 mm) were cleaned first with 100% (v/v) isopropanol followed by pure MilliQ water. The procedure was repeated twice and then slides were air dried using a pressurized air stream. Silicon gaskets to hold the sample drops were cleaned in the same manner as described for the slides. All preparation were done prior to immediate start of the experiment. All data were recorded using AcquireMP software. The instrument was calibrated using NativeMark protein standards in the same sample buffer (10 mM Tris pH 9.0, 200 mM NaCl, 5 mM DTT) prior to sample measurements. Stock protein sample was diluted to final concentration between 100 nM concentration. To find focus 8uls of buffer was pipetted into a well and the focal position was identified and locked using the autofocus function. Then to acquire sample measurements 2 μL of 100 nM concentration of protein was pipetted into 8 μL of the buffer and mixed carefully. Mass photometry movies were then recorded for 120 seconds. The data were analysed using the DiscoverMP software and generated mass distribution histograms.

### Cryo-electron microscopy (Cryo-EM) for determination of *Ndas*CDO structure

*Sample preparation for cryoEM* – Prior to preparing grids, the purified *Ndas*CDO complex dialysed against 20 mM Tris pH 9, 200 mM NaCl, 5 mM DTT for 2-3 hrs. The dialysed protein was diluted to 4 μM final concentration and spun at 8000 rpm on a Eppendorf table top centrifuge for 5 minutes to remove any aggregates. Grids of the NDS protein were then prepared by applying 4 μL of 4 μM protein to a glow-discharged (1 min at 0.1 atmospheric pressure, 35 mAmps, Quorum technologies SC7620) R2/1 400 mesh Cu/Rh holey carbon grids. The grids were then vitrified by plunging into liquid ethane using FEI Vitrobot Mark IV. Before plunging, grids were blotted for 3 seconds with the blot force of 3.5, and the climate chamber was maintained at 4 °C and 100 % humidity. The grids were subsequently clipped and stored in the liquid nitrogen until data collection.

*Data collection, processing, and model building –* Initial screening of *Ndas*CDO grids were carried out in our in-house Glacios microscope equipped with Falcon4i camera (University of Dundee, CryoEM facility). Best grids with well-defined ice and good distribution of filaments are then transported to eBIC cryo electron microscopy facility located at Harwell Diamond. Data were then collected using a 300Kv Titan Krios cryo electron microscope equipped with Gatan K3 direct electron detector. EPU software was used to select targets and acquire movies in super resolution electron counting mode at a magnification of 105kx and calibrated sampling pixel size of 0.831 Å/pixel. In total 12900 movies were collected with 50 frames per movie stack and exposure time of 1.79 seconds per movie stack using aberration free image shift (AFIS).

Structural reconstruction using helical processing methodology was carried out using cryoSPARC suite (v4.4.1)^27^. Movies were imported into Cryosparc and using patch motion correction frames are aligned and beam induced motions correction, and electron dose weighting were performed. CTF estimation using CTFFIND4 was carried out and those micrographs with CTF fit worse than 4.8 Å were discarded which results in 9031 images. Particles were initially picked and optimised on a subset of 200 images using Filament tracer with filament diameter of 120 Å which is subsequently extended to all the images. A total of 1.8 M particles were picked and extracted with a box size of 340 pixels. The extracted particles were then inspected, filtered for any ice contamination and artifacts, and subjected to three rounds of 2D classification. The best 2D classes were selected, resulting in 849,600 particles and a proportion of particles (180 k) from the classes were used for initial Ab-initio model generation. Subsequently all the 849 k particles and the ab-initio model as input for a Helix Refine job. The map generated from the Helix Refine job had a global resolution of 3.6 Å resolution with the estimated helical twist of 132.3° and a helical rise of 45.45 Å. The particle stack was then subjected to Global and local per particle CTF refinement before performing second round of Helical refinement which improved the map to 3.4 Å resolution. A local refinement was carried out using a mask covering two dimers of *Ndas*CDO subunit A and two dimers of *Ndas*CDO subunit B, yielding a final map of 3.07Å global resolution. The final map was filtered and sharpened using B-factor of 180 Å^2^.

AlphaFold2^28^ was used to generate an initial model of the individual *Ndas*CDO subunit A and B monomers. The models were then rigid body docked into the filament map and respective oligomers were generated. The structure was refined with rounds of model building in Coot, fitting with adaptive distance restraints in ISOLDE (v1.6)^29^ and refinement with Phenix (v1.20.1-4487)^30^ real-space refinement. Figures were generated in ChimeraX (v1.6)^31^ and The PyMOL Molecular Graphics System, version 1.8 (Schrodinger, 2015).

### Electron Paramagnetic Resonance (EPR) Spectroscopy

5 mM cFG or 5 mM cGG were added to 100 μM *Ndas*CDO and each reaction mixture was frozen using liquid nitrogen. Controls containing only 5 mM cFG or 100 μM NdasCDO were also prepared. Continuous wave (CW) EPR spectra were obtained at 120 K with a Bruker EMX 10/12 spectrometer running Xenon software and equipped with an ELEXSYS Super Hi-Q resonator at an operating frequency of ∼9.50 GHz with 100 kHz modulation. Temperature was controlled with an ER4141 VTM Nitrogen VT unit (Bruker) operated with liquid nitrogen. CW spectra were recorded using a 20 mT field sweep centred at 336 mT, a time constant of 40.96 ms, a conversion time of 20 ms, and 1000 points resolution. An attenuation of 23.0 dB (1 mW power) and a modulation amplitude of 0.2 mT were used. CW spectra were phase– and background-corrected using the Xenon software.

### pH-rate profiles

Prior to the execution of pH-rate profiles, enzymatic stability tests were performed. *Ndas*CDO was diluted to 10 μM in H_2_O or 200 mM of a mixed buffer containing MES, Tris, CAPS and KCl at either pH 6 or 11. *Ndas*CDO, at a final concentration of 0.1 μM, was then added to a reaction mixture containing 50 mM Tris pH 8 and 200 mM NaCl. cFP was used at saturating concentrations to verify that *NdasCDO* retained activity following incubation at both low and high pH.

For pH profiles, the mixed buffer system was prepared to range from pH 6-11 in increments of 0.5 pH units. A final concentration of 200 mM of each buffer was used whilst cFP substrate concentration was varied. *Ndas*CDO was added last to a final concentration of 0.1 μM and all reactions were measured in triplicate.

### Temperature-rate profiles

Firstly, to determine whether *Ndas*CDO was stable at the temperatures used in this study, *Ndas*CDO was incubated at the extremes of the experimental temperature range and subsequently assayed at 298 K. This confirmed no loss of enzymatic activity occurring upon incubation for 30 minutes.

For temperature investigations, cFP was used at saturating conditions in 50 mM Tris, pH 9 for optimal activity as confirmed by the previous pH-rate profile. *Ndas*CDO was added to a final concentration of 0.1 μM and the activity was measured between a temperature range of 298 to 318 K in 5 K increments, which was dictated by the constraints of the plate reader used.

### Solvent Kinetic Isotope Effects (SKIEs)

Solvent kinetic isotope effects (SKIEs) were explored by plotting saturation curves in H_2_O or 91% D_2_O. Additionally, a control experiment containing 9% glycerol was performed concurrently. Here, cFG was used as a substrate with 0.1 μM *Ndas*CDO in 50 mM Tris pH 9 and 200 mM NaCl. For proton inventory experiments, D_2_O was varied from 0 to 90% in increments of 10%. Buffers prepared in D_2_O were corrected for pH alterations caused by the presence of D_2_O [pD+ = pHa (apparent reading from pH meter) + 0.4]. SKIE investigations in H_2_O were analysed using Michaelis-Menten analysis to obtain both *k*_cat_ and *K*_M_.

### Viscosity studies

Potential viscosity effects were investigated using either sucrose or glycerol as the viscogen. Relative viscosities (η_rel_) for different concentrations of glycerol and sucrose were measured by Bazelyansky *et al.* and listed in the supporting information on Table S2.

For viscosity effects, cFG was used as the substrate at varying concentrations with 0.1 μM of *Ndas*CDO in 50 mM Tris pH 9 and 1 M NaCl. Each experiment was incubated at 30 °C in triplicate and the absorbance was measured at 297 nm.

### Determination of extinction coefficients

Spectrophotometric assays were used to monitor the oxidation of cyclodipeptides. Initially, the absorbance spectrum (200 – 1000 nm) was measured upon the addition of *Ndas*CDO (i.e., 0 minute timepoint) and then again after 24 hour incubation at 30 °C. From here, the difference in absorbance for each respective oxidised CDP was used to create a progress curve. The last 20 points for each concentration were averaged and plotted against the substrate concentration to give a linear trend where the gradient represented the extinction coefficient for the oxidised CDP in absorbance/concentration (M) units. Figure S2 depicts UV difference spectra upon oxidation for different CDP substrates.

### LC-MS Analysis of oxidised cyclodipeptides

Samples containing 1 μM *Ndas*CDO and 30 μM of the respective cyclodipeptide were incubated in 50 mM Tris pH 8 and 200 mM NaCl overnight at room temperature. The reaction was quenched upon the addition of cold methanol to a final concentration of 80%. The samples were incubated at –80 °C for 15 minutes and centrifuged at 18000 x g for 10 minutes. The resultant supernatant was dried under nitrogen gas and reconstituted in LC-MS grade water. Raw data is available on Figure S3a-h.

All samples except for cWS were analysed using a Waters ACQUITY UPLC liquid chromatography system coupled to a Xevo G2-XS Qtof mass spectrometer equipped with an electrospray ionization (ESI) source. 1 μL of sample was injected onto the appropriate column and ran at 40 °C. All histidine or tryptophan containing compounds were injected onto a HSS-T3 column (2.1 x 100 mm, 1.8 μm, Waters Acquity) whilst non-polar CDPs such as cFP or cLP were run on a BEH C18 column (2.1 x 100 mm, 1.7 μm, Waters Acquity). CDPs were separated from the mixture using a gradient mobile phase from 1% B to 50% B where the two mobile phases consisted of A – 0.1% formic acid in water and B – 0.1% formic acid in acetonitrile at a flow rate of 400 μL min^−1^. The capillary voltage was set at 2.5 kV in positive ion mode. The source and desolvation gas temperatures of the mass spectrometer were set at 120 °C and 500 °C respectively. The cone gas flow was set to 50 L/hr whilst the desolvation gas flow was set at 1000 L/hr. An MSE scan was performed between 50 – 700 m/z where function 1 employed MS analysis whilst function 2 applied a collision energy ramp from 15 to 30 V to perform MS/MS fragmentation. In addition, a lockspray signal was measured and a mass correction was applied by collecting every 10 s, averaging 3 scans of 1 s each using 50 pg mL^−1^ of Leucine Enkephalin dissolved in water:acetonitrile (50:50) and 0.1% formic acid as a standard (556.2771 m/z, Waters).

For cWS, samples were subjected to LCMS/MS using an Eksigent Ekspert nanoLC 425 (Eksigent, AB SCIEX) coupled to an Triple ToF 6600 mass spectrometer (ABSCIEX). Dipeptides were injected on a reverse-phase YMC Triart C18 trap column (12nm, 3 μM, 0.3 x 0.5 mm) for pre-concentration and desalted with loading buffer at a flow rate of 5ul/min for 1min. The peptide trap was then switched into line with a YMC Triart C18 analytical column (12 nm 3 μm 0.3 x 150 mm) column. Peptides were eluted from the column using a linear solvent gradient using the following gradient: linear 3–95 % of buffer B within 6min, isocratic 95 % of buffer B for 2 min, decrease to 3 % buffer B within 1min and isocratic 2 % buffer B for 4 min. The mass spectrometer was operated in positive ion mode and acquired a TOF MS scan for 120 msec of accumulation time from m/z 80-500, followed by three Product Ion scans for the respective reaction product with the same accumulation time and m/z scan range with a cycle time of 0.5 seconds.

### Pre-steady state multiple turnover

Concentrations reported are final after mixing 30 μL each sample in a 1:1 ratio in an Applied Photophysics SX20 stopped flow instrument, with a cell of 5 μL, a pathlength of 0.5 cm and with an instrument deadtime of 1 ms. Multiple turnover experiments were carried out as follows:

*cFP:* 8.5 μM *Ndas*CDO and varying concentrations of the cFP (0.5, 1 and 2 mM) in 50 mM Tris pH 9 and 200 mM NaCl. Reaction was monitored at 312 nm for 5 s, 10000 datapoints were collected in logarithmic scale. Voltage applied was 384.4 V.

*cWS*: 8.5 μM *Ndas*CDO and varying concentrations of cWS (0.5 and 1 mM) in 50 mM Tris pH 9 and 200 mM NaCl. Reaction was monitored at 297 nm for 5 s, 10000 datapoints were collected in logarithmic scale. Voltage applied was 400 V.

Data were fitted both analytically using Graphpad Prism (exponential equations in the “Data fitting” section below and numerically using Kintek Global Explorer^32^.

### Mass spectrometric investigation of flavin cofactor

*Trypsin digestion sample preparation –* Protein was digested into peptides by adding trypsin (1:50 w/w protease:protein) and incubating over night at 30C.

*Data acquisition –* Peptides were subjected to LCMS/MS using an Ultimate 3000 RSLC (Thermo Fisher Scientific) coupled to an Orbitrap Fusion Lumos mass spectrometer (Thermo Fisher Scientific) and equipped with FAIMS. Peptides were injected onto a reverse-phase trap (Pepmap100 C18 5 μm 0.3 × 5 mm) for pre-concentration and desalted with loading buffer, at 15 μL/min for 3 minutes. The peptide trap was then switched into line with the analytical column (Easy-spray Pepmap RSLC C18 2μm, 15 cm x75 μm ID). Peptides were eluted from the column using a linear solvent gradient using the following gradient: linear 4–40 % of buffer B over 45 min, linear 40–95 % of buffer B for 4 min, isocratic 95 % of buffer B for 6 min, sharp decrease to 2 % buffer B within 0.1min and isocratic 2 % buffer B for 10 min. The FAIMS interface alternated between –45 and –65V The mass spectrometer was operated in DDA positive ion mode with a cycle time of 1.5 sec. The Orbitrap was selected as the MS1 detector at a resolution of 60000 with a scan range of from m/z 375 to 1500. Peptides with charge states 2 to 5 were selected for fragmentation in the ion trap using HCD as collision energy. The raw data files were converted into mgf using MSconvert (ProteoWizard) and searched using Mascot with trypsin as the cleavage enzyme and Flavin mononucleotide FMN [C(17)H(21)N(4)O(9)P with a delta mass of 456.1046] as a variable modification of cysteines as well as oxidation as a variable modification of methionines against an in-house database containing 42354 protein sequences. The mass accuracy for the MS scan was set to 20ppm and for the fragment ion mass to 0.6 Da. Trypsin was selected as the enzyme with 1 missed cleavage allowed.

*Intact protein mass analysis –* The protein sample (20μL, 1 μM) was desalted on-line through a MassPrep On-Line Desalting Cartridge 2.1 x 10 mm, using a Waters Acquity H-class HPLC, eluting at 200 μL/min, with an increasing acetonitrile concentration (2 % acetonitrile, 98 % aqueous 1 % formic acid to 98 % acetonitrile, 2 % aqueous 1 % formic acid) and delivered to a Waters Xevo G2XS electrospray ionisation mass spectrometer operated with positive polarity in sensitivity mode. Intermittently a lockspray signal using Leucine Enkephalin was measured and a mass correction was applied. An envelope of multiply charged signals was acquired between m/z 500-2500 and deconvoluted using MaxEnt1 software to give the molecular mass of the protein.

## Data fitting

The entire range of the temperature were fitted to an Eyring equation (Equation 1) ^33–35^, where R is the gas constant; *T* is the temperature (K); T_0_ is a reference temperature value (30 °C or 303.15 K was used); *k* is the kinetic rate constant or parameter; Δ*H*^‡^ is the enthalpy of activation; *k*_B_ the Boltzmann constant; ℎ the Planck’s constant; Δ*S*^‡^ the entropy of activation.

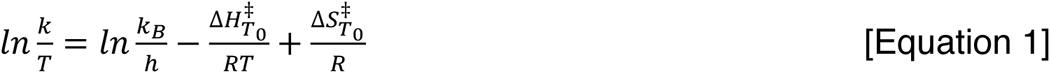

pH data were plotted with pH and log kinetic parameter. Data for *k*_cat_ pH-rate profiles were fitted to Equation 2 (accounting for one ionizable group in each limb), while data for *k*_cat_/*K*_M_ were fitted to Equation 3 (accounting for two ionizable groups in each limb). For Equations 2 and 3, C is the pH independent value of the kinetic parameter; p*K*_a1_ and p*K*_a2_ are the dissociation constants for ionizable groups. Equation S1 is an alternative to when ionizable groups are close together, and its usage (or not) is discussed in the Supporting information.

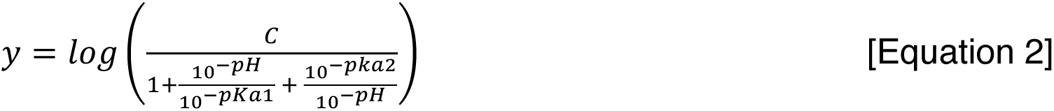

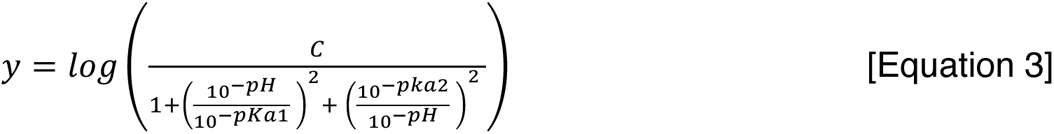

When calculating SKIEs, *k*_cat_ and *K*_M_ were obtained from the Michaelis-Menten curves in H_2_O. The 91% D_2_O Michaelis Menten curve was fit to the following equation:

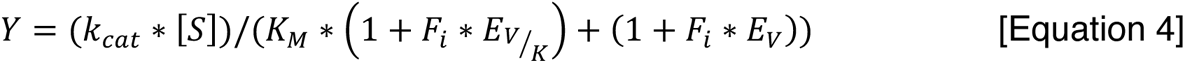

Where F_i_ is the fraction D_2_O, and E_V/K_ and E_V_ are the isotope effects –1 on *k*_cat_/*K*_M_ and *k*_cat_ respectively.

Fitted values for kinetic parameters were converted into energy barriers using Equation 5:

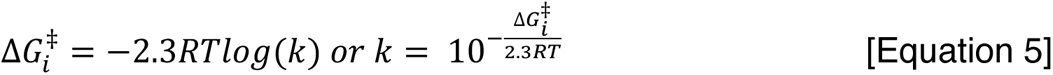

Differential scanning fluorimetry data were fitted to the standard Boltzmann sigmoidal function where Bottom refers to the lowest value, Top is the largest and V50 is the median of those two values:

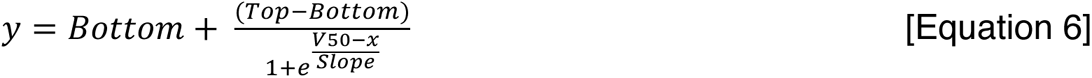

Pre-steady state multiple turnover data were fitted to a standard exponential equation followed by a linear slope:

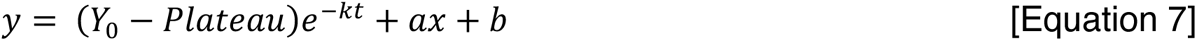

For equation 7, t is time in seconds, Y_0_ is the lowest value of Y, Plateau is the Y value in which the exponential phase ends, *k* is the rate constant in s^−1^, a is the slope of the linear phase and b is the y intercept from the linear phase.

For data fitting using Kintek Global Explorer^32^, the following Model was used, in which CDP is either cFP or cWS:

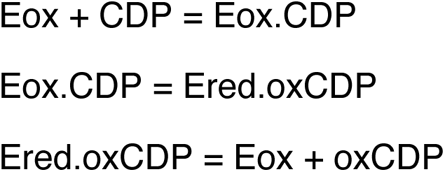

The signal observed was set as below, where scale_1a is a concentration scaling factor, a is the fluorescence contribution for the species being detected, Ered.oxCDP is the *Ndas*CDO-oxidized product complex and oxCDP is the oxidized product, b is the background:

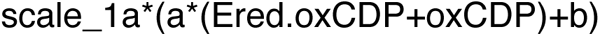

## Results and Discussion

### General characterization of *Ndas*CDO

*Covalent FMN cofactor* – *Ndas*CDO purifies with a yellow colour and characteristic absorption spectra for flavin-dependent enzymes (Figure S4). Protein intact mass spectrometry shows a mass shift of + 907 Da for the A subunit dimer (Figure S2), indicative of one FMN cofactor bound per subunit. Trypsin digestion and peptide fingerprinting confirmed the assignment of Cys121 as the covalent attachment site for FMN. Prior work with CDOs under anaerobic conditions observed no cyclodipeptide oxidation^20^, and therefore the catalytic cycle starts with the cofactor as the oxidized quinone form of the cofactor (Figure 1a, quinone; Fl_ox_). Covalently attached flavin cofactors are observed in approximately 10% of flavoenzymes, and this attachment has been proposed to modulate cofactor redox potential, therefore facilitating catalysis of thermodynamically challenging reactions.^36, 37^

*Temperature-rate profiles and stability* – *Nocardiopsis dassonvillei* is a halophilic bacterium, with optimal growth conditions containing up to 3.4 M NaCl^38^, therefore we explored whether high salt conditions were required for protein stability and activity. Folding and melting temperatures of *Ndas*CDO revealed a Tm of 46 °C with 0.2 M NaCl, and that higher salt concentrations were associated with higher melting temperatures (Figure 1a). Next, we analysed the temperature dependence of cFP oxidation catalyzed by *Ndas*CDO (individual Michaelis-Menten curves available on Figure S5a). Figure 1c depicts a slight curvature in the temperature range under evaluation. However, we opted for a simple model using a linear fit to the Eyring Equation without a contribution from activation heat capacity, further discussed below, with fitted values for the temperature dependence of *k*_cat_ of Δ*H*^‡^ = 4.06 ± 2.42 kcal mol^−1^ and Δ*S*^‡^ of –0.049 ± 0.008 kcal mol^−1^. In our standard reaction conditions (30 °C), a Δ*G*^‡^ = 16.5 kcal mol^−1^ would be observed, mostly driven by entropic contributions. The relevance of entropic contributions in enzyme catalysis has been discussed in the context of the importance of interactions between solvent and outer protein surface^39^, and whether the large complex formed by subunits A and B plays a role remains to be determined.

Care must be taken when evaluating curved temperature profiles in enzyme-catalyzed reactions, as many reasons for curvature exist (such as changes in rate limiting step, contributions of heat capacity, compensatory effects from contributions of enthalpy and entropy in the range under investigation).^34^ Figure S5b depicts a comparison between fits to an Eyring Equation with and without contributions from activation heat capacity, as well as with linear fits that would occur due to changes in the rate limiting step for the reaction under study. Because we have no additional experimental evidence to invoke a model accounting for curvature, compounded with the fact that the filamentous structure of *Ndas*CDO could further complicate data interpretation, we therefore report a linear dependence as the best fitted model.

*pH dependence* – *Ndas*CDO depicted bell shaped profiles for both *k*_cat_ and *k*_cat_/*K*_M_ (Figure 1b). Best fitted values were obtained with a single ionizable group on each side of the *k*_cat_ profile (p*K*_a1_ = 8.3 ± 0.1, pK_a2_ = 9.8 ± 0.1), but with two ionizable groups on each side of the *k*_cat_/*K*_M-cFP_ profile (p*K*_a1_ = 8.6 ± 0.1, pK_a2_ = 10.0 ± 0.1). These experiments were performed with cFP as a substrate, and therefore no contribution from substrate ionizable groups is expected. Structural data led us to hypothesize one of these ionizable groups observed as contributing to pK_a1_ could be Y36 (subunit B). pK_a2_ could be reporting on the FMN cofactor, since no other ionizable groups with predicted pK_a_ values > 8.0 are present in the vicinity of FMN. A proposed pK_a_ of ∼8.3 for the flavin semiquinone in vanillyl-alcohol oxidase has been reported^40^, and future studies on the precise nature of this ionizable group as well as flavin intermediates formed during turnover can be performed to better ascertain group(s) giving rise to the pH-rate profiles observed here.

After pH and temperature studies, the best reaction conditions to monitor the *Ndas*CDO-catalyzed reaction were established (30 °C at 50 mM Tris, pH 9.0 with 200 mM NaCl), and then employed for subsequent substrate scope, viscosity studies, solvent kinetic isotope effects and proton inventory studies (discussed below).

*Oligomeric state* – Purified *Ndas*CDO displayed a large molecular weight, as previously observed for other CDO for which a preliminary characterization was performed^20, 22^. We performed mass photometry experiments (Figure 1f) to determine a salt-dependent oligomerization process, in which higher salt concentrations led to a more homogeneously distributed population of *Ndas*CDO with a MW of 205 ± 47 kDa. Functional enzyme contains subunit A (22kDa) and subunit B (11kDa) at an unknown ratio. These results suggest *Ndas*CDO is a filament, as observed in a homologue enzyme^41^.

### *Ndas*CDO is a filament enzyme

*Atomic structure of NdasCDO filaments* – To gain insights into the structural organisation of *Ndas*CDO, we performed helical reconstruction from the cryo electron microscopy data collected from a cryo vitrified grid containing ordered *Ndas*CDO filaments (Figure 2a and b) to resolve a 3.0 Å resolution map. This yielded a high quality map, to which the atomic model containing both A and b subunits was fitted. This model was generated using the alphafold model for the individual monomers of subunit A and B (Figure 2c, Figure S6, Figure S7). Figure S8 shows the data processing pipeline used.

**Figure 2:**
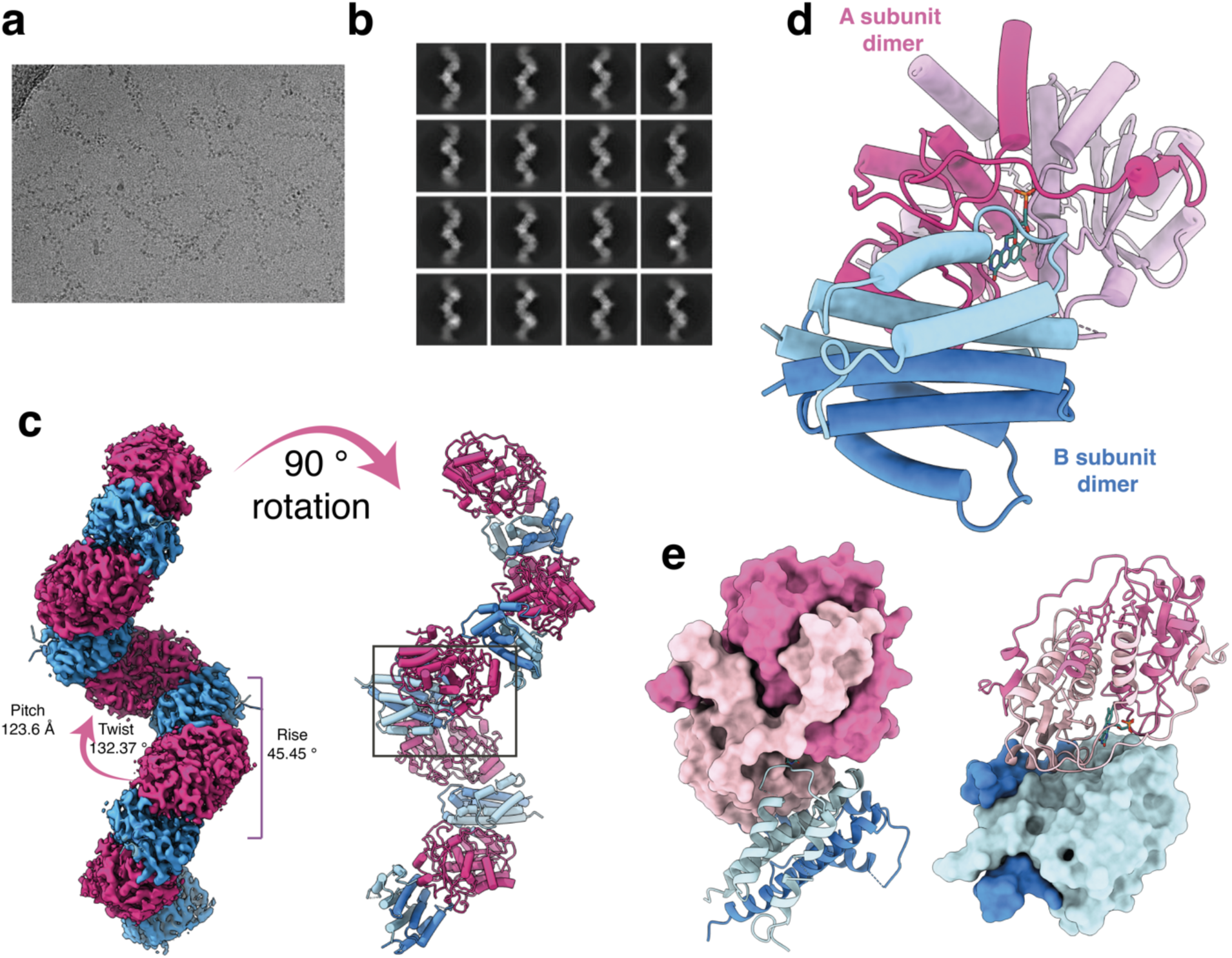
*Ndas*CDO filament structure revealed by cryo-EM. For all panels, A subunits are shown in shades of pink and B subunits in shade of blue. a) Data acquired showing particles and b) representative image depicting best 2D particles picked for map generation. c) Cryo-EM density resulting from the helical reconstruction of the *Ndas*CDO filament. On the right side, filament is rotated and subunits shown as cartoons. Box highlights the functional dimer or dimers between A and B subunits. d) View of the functional dimer or dimers between A and B subunits. The FMN cofactor (teal, sticks) is flanked by residues from both A subunits and one of the B subunits. e) Surface representation of the refined models for subunits A (left) and B (right).

The filament is composed of alternating repeating units of homodimers of *Ndas*CDO subunit A (*Ndas*CDO-A) and homodimers of *Ndas*CDO subunit B (*Ndas*CDO-B), such that the interface between the two homodimers constitutes the heterotetrametric form of the molecule representing the biologically relevant asymmetric unit. Continuous filament is formed by shifting the homo-tetramer along the screw axis with helical rise of 45.45 Å, helical twist of 132.37° and overall helical pitch of 123.6 Å (Figure 2c).

*Subunit A*: The structural fold of *Ndas*CDO-A resembles that of the flavin-dependent nitro reductase family of proteins^42^. *Ndas*CDO-A is composed of four anti-parallel beta strands surrounded by five alpha helical elements (Figure 2d and Figure 2e). The largest alpha helix α4 (Figure S7a) lies parallel to the β-sheet carries the highly conserved cysteine Cys121. Within the *Ndas*CDO-A map, clearly evident compatible density is found for the FMN cofactor stemming from the Cys121 aa. The modelled FMN cofactor clearly fits the density whose 8α carbon lies at distance of ∼1.7 Å to the thiol group of Cys121 confirming the covalent linkage with FMN (Figure S7d). The α4 helix along with covalently linked FMN plays central interaction site for the homo-dimer formation (Figure S7b). Surrounding the α4 helix two other alpha helices α1 and α2 from each monomer stack against the α4 and together form the central helical core interaction of the dimer. The C-terminal region from β4 (region 172-195) forms an extended region, carrying a small 3_10_ helix and a β-strand that sits parallel to the β1 of the second monomer. Altogether the intact homo dimer forms the active site region for the FMN cofactor and potential binds the CDP substrate.

*Subunit B*: *Ndas*CDO-B is an α-helical protein composed of 4 antiparallel helices and together with the second monomer forms a dimeric 8-helical bundle structure (Figure 2d, Figure 2e, and Figure S7b). Residues from both the monomer carrying a heptad repeat element pattern and through extensive inter-molecular interaction forms the interlock between the two monomer helices and creates a knob-into hole structure. Helices α1, α2 and α3 from one monomer together with α4 from second monomer creates hetero tetramer interface with a subunit from the *Ndas*CDO-A homodimer.

*Interactions with the FMN cofactor:* The FMN cofactor is covalently attached to Cys121 and sandwiched between subunits A and B. Covalent attachment to subunit A and to this cysteine residue specifically was confirmed by protein mass spectrometry (Figure S4). R28, R29 and R181 participate in polar interactions with the phosphate group of FMN, R32 is in close proximity to one of the C2 carbonyl from FMN’s isoalloxazine ring (Figure 3a and Figure S7). However, variants harbouring a truncated N-terminus (NdasCDO_△2-17_ and NdasCDO_△2-39_), specially *Ndas*CDO_△2-39_ which removed R28, R29 and R32 retain catalytic activity. *Ndas*CDO_△2-17_ is comparable to wild type in the formation of singly and doubly oxidized products, and NdasCDO_△2-39_ can catalyze CDP oxidation, albeit with much lower yields. It is likely that this truncation impairs catalytic turnover, given polar contacts to optimally position the FMN cofactor. Nevertheless, these truncations demonstrate that the flexible N-terminus of CDO-A is not required for function.

**Figure 3:**
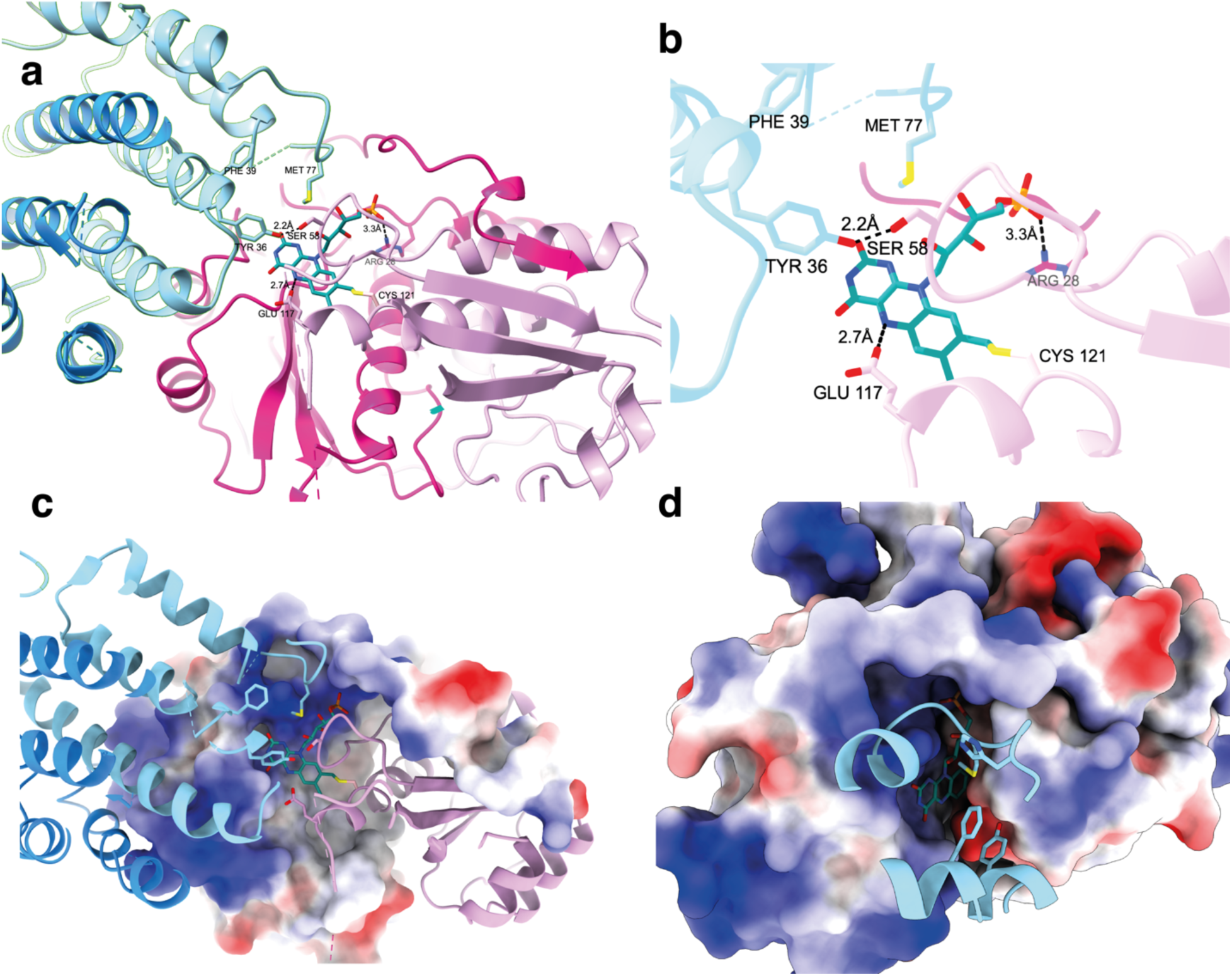
FMN binding site and active site environment. a) Overview of *Ndas*CDO-A (pink) and *Ndas*CDO-B (blue) subunits, FMN (teal) is covalently linked to Cys121, and residues in the vicinity of the cofactor are shown. Figure S7d depicts continuous density between Cys121 and the FMN cofactor. b) Key residues interacting the with FMN cofactor are shown. One monomer from B subunit covers the active site, and Met77, Phe39, Tyr36 are shown as sticks. From *Ndas*CDO-A, Glu117 is in close proximity to the N5 of FMN, Ser58 is in close distance to *Ndas*CDO-B Tyr36. From a second *Ndas*CDO-A monomer (dark pink), Arg 28 is in a polar interaction with the phosphate group of the cofactor. c) One monomer of *Ndas*CDO-A is shown as surface, colored by electrostactics. The putative substrate binding pocket is therefore covered by Met77 and Tyr36 from *Ndas*CDO-B. d) Both monomers of *Ndas*CDO-A are shown as surface, colored by electrostactics, and key regions from *Ndas*CDO-B are shown as cartoons. FMN is in a deep pocket, mostly surrounded by positively charged and hydrophobic residues. For all panels, FMN is in teal, one subunit A is in dark pink and the other on light pink, one B subunit is shown in light blue, and the other in darker blue.

A DALI search revealed the closest homologue in the pdb to *Ndas*CDO-A to be the nitroreductase NfsA^42^ (Figure S10). Catalytic Ser48 of NfsA is crucial for function but mutating the equivalent serine residue to alanine (*Ndas*CDO-A_S58A_) does not lead to complete loss in function (Figure S11). Nevertheless, *Ndas*CDO-A_S58A_ has impaired *k*_cat_ and *k*_cat_/*K*_M_, and given the proximity to Tyr36-B (*Ndas*CDO-B), it could be important in the catalytic mechanism of the reaction. Glu117 from one of the monomers of *Ndas*CDO-A is within hydrogen bonding distance from the N5 of FMN. Glu117 is 7 Å from Ser58-A or Tyr36-B, too far for a direct interaction but nevertheless in a position that could lead to interactions with CDP substrates and their side chains are participate in proton transfer with the cofactor. Together Tyr36-B, Phe39-B and Met77-B can limit access to the substrate binding site and FMN, as they cover the active site (Figure 3d). The FMN and substrate binding pocket are in a deep pocket, mostly surrounded by positively charged and hydrophobic residues (Figure S7), and further shielded from solvent by Tyr36-B, Phe39-B and Met77-B (Figure 3c and Figure 3d).

### Substrate scope of *Ndas*CDO

*Substrate scope* – *Ndas*CDO is a promiscuous enzyme, accepting multiple CDP substrates^22, 25^ (Figure 4). Assay conditions and procedure to determine extinction coefficients employed are available in the supporting information (Figure S2). Table 1 summarizes kinetic parameters for distinct CDP substrates. *K*_M_ values were between 0.08 mM (cWF) and 4.2 mM (cFG), indicating overall poor affinity for CDP substrates. Surprisingly, cHF was the best substrate (*k*_cat_/*K*_M-cHF_ = 30.6 ± 0.1), mostly due to a *k*_cat_ effect. The CDPS enzyme part of the same biosynthetic gene cluster *Ndas*CDO is a natural producer of cWF, a substrate with overall poor efficiency, and in the same magnitude of cLP, demonstrating a non-strict preference for aromatic amino acid side chains.

**Figure 4:**
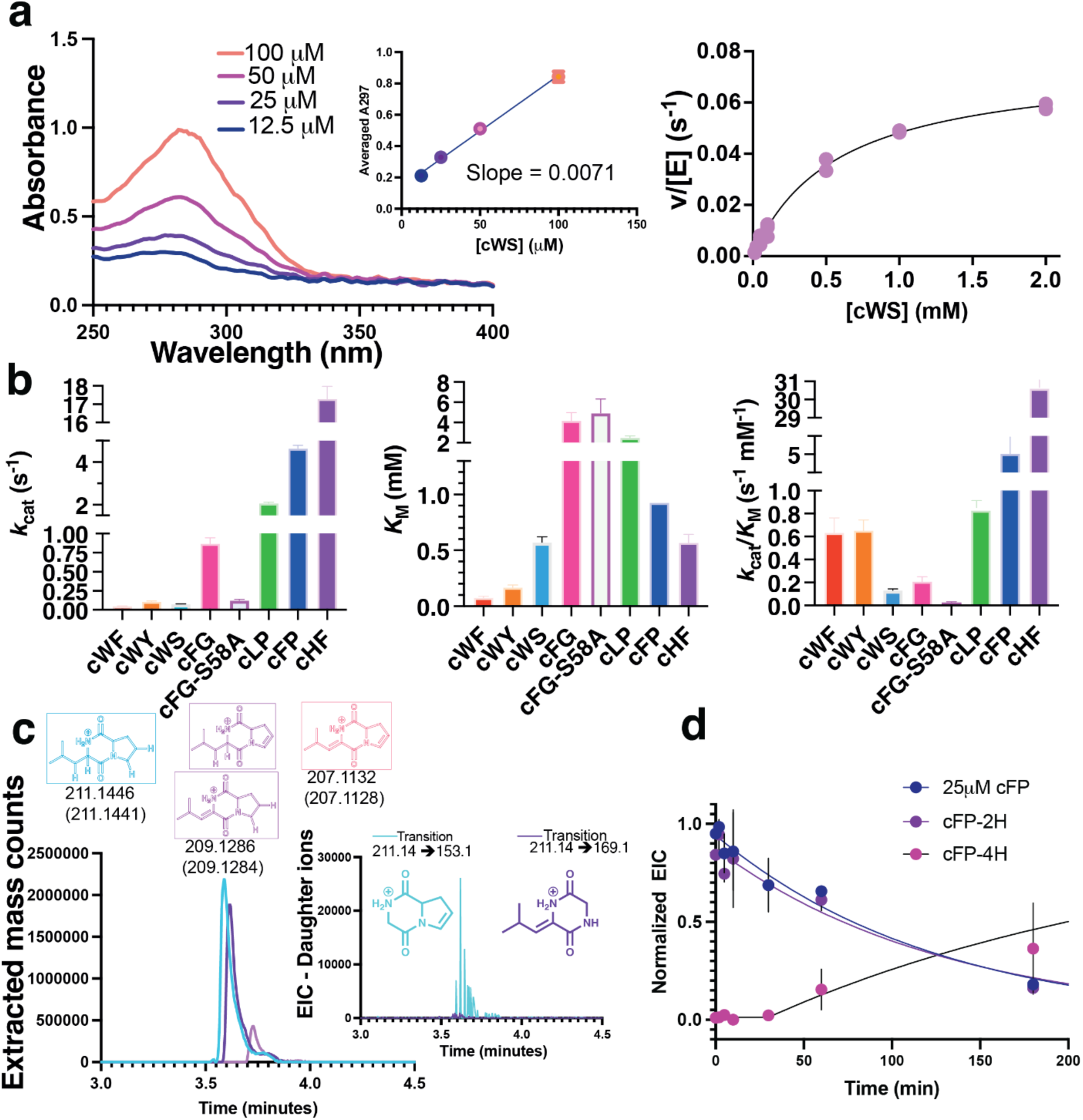
Substrate scope of *Ndas*CDO. a) overview of workflow to test different CDP substrates and determine kinetic parameters. First (left), progress curves were acquired overnight, and after reaction had reached completion final absorbance readings were utilized to generate a calibration curve correlating change in absorbance to amount of product formed (middle). Then, assays under initial rate conditions were carried out to determine substrate scope and preference (right). Only absorbance readings including under 10% substrate conversion into products were considered. b) Fitted values for *k*_cat_ (left), *K*_M_ (middle), and *k*_cat_/*K*_M_ (right) are shown with standard deviation. All experiments were carried out in triplicate and raw data for individual progress curves are available on Figure S2, all Michaelis-Menten plots available on Figure S12. c) Unique transitions observed by mass fragmentation of cLP-2H oxidation (Figure S3i). Inset shows in blue a unique transition if cFP-2H underwent dehydrogenation on the Pro sidechain, while purple (not present) depicts unique transition if cFP-2H underwent dehydrogenation on the Phe sidechain. Fragmentation was determined as previously described^25^. d) Reaction with cFP monitored by mass detection, using single reaction monitoring (SIR) channels for substrate, product with 1 oxidation (cFP-2H) or 2 oxidations (cFP-4H). Under these conditions there is a lag for formation of cFP-4H, indicating sufficient levels of cFP-2H needed to accumulate before a second dehydrogenation reaction could take place (Figure S13).

**Table 1:** Kinetic parameters for *Ndas*CDO substrates.

| | $k_{cat}$ (s<sup>-1</sup>) | $K_M$ (mM) | $k_{cat}/K_M$ (mM<sup>-1</sup> s<sup>-1</sup>) |
| --- | --- | --- | --- |
| **cWF** | 0.05 ± 0.01 | 0.08 ± 0.01 | 0.63 ± 0.21 |
| **cWY** | 0.11 ± 0.01 | 0.17 ± 0.02 | 0.65 ± 0.15 |
| **cWS** | 0.076 ± 0.002 | 0.57 ± 0.05 | 0.13 ± 0.01 |
| **cFG** | 0.87 ± 0.08 | 4.2 ± 0.8 | 0.21 ± 0.21 |
| **cFG-S58A** | 0.12 ± 0.02 | 4.9 ± 1.4 | 0.025 ± 0.008 |
| **cLP** | 2.07 ± 0.06 | 2.5 ± 0.2 | 0.83 ± 0.07 |
| **cFP** | 4.6 ± 0.2 | 0.9 ± 0.1 | 5.0 ± 0.1 |
| **cHF** | 17.3 ± 0.7 | 0.57 ± 0.08 | 30.6 ± 0.1 |

**Table 2:** Rate constants obtained after fitting multiple turnover experiment.

|  | **Best fitted value** | **Lower boundary\*** | **Upper boundary** |
| --- | --- | --- | --- |
| $k_1$ | $0.0031 \pm 0.0001$ | 0.0025 | 0.0038 |
| $k_{-1}$ | $16.8 \pm 0.8$ | 18.2 | 38.6 |
| $k_2$ | $32.0 \pm 0.5$ | 49.2 | 75.3 |
| $k_{-2}$ | $1.8 \pm 0.1$ | 1.6 | 2.2 |
| $k_3$ | $2.0 \pm 0.1$ | 1.8 | 1.8 |

\* Boundaries were calculated with $\chi^2 = 0.95$ , and $k_{-3}$ was fixed at zero, since product release was considered irreversible under multiple turnover conditions.

Other CDPs were tested as substrates (cWG, cWW, cWS, cFF, cHP), and were successfully oxidized (verified by LC-MS, Figure S3), however a combination of high *K*_M_ values, slow reactions, substrate solubility and small changes in absorbance due to product formation did not allow determination of kinetic parameters for all substrates tested. LC-MS assays revealed cWG underwent slow oxidation with little product formed after 2h reaction.

We tested cWS as a substrate as the acceptance of substrates with polar side chains was undetermined. Furthermore, cWS is an interesting peptide with antibiofilm activity against *Pseudomonas aeruginosa*.^43^ With purified reaction components, *Ndas*CDO can catalyze single and double dehydrogenation, although the product cWS-2H accumulated after an overnight reaction, indicative that the second dehydrogenation was not favoured. Prior work with cell cultures and *Ndas*CDO failed to observe cWS as a feasible substrate^22^, and we hypothesize this could be due to 1-the high *K*_M_ value for cWS (0.8 mM), and 2-the higher complexity and suppression effects observed when carrying out LC-MS assays with complex mixtures for CDP detection^44^.

*Studies on second oxidation* – Most substrates tested can be doubly oxidized by *Ndas*CDO (Figure S2). We performed a time course experiment monitoring by mass detection the appearance of the first oxidation product followed by the second oxidation product. Based on progress curves (Figure 4d) monitoring cFP disappearance and appearance of cFP-2H and cFP-4H (for products with one or two oxidized bonds, respectively). cFP is completely converted into cFP-4H in an overnight reaction. We propose *Ndas*CDO is releasing a CDP-2H product, which then must rebind in a different orientation so that second oxidation can occur. Overall, substrates containing one tryptophan were poor candidates for formation of doubly-oxidized products, while substrates containing Phe and other small hydrophobic side chains were more efficient substrates.

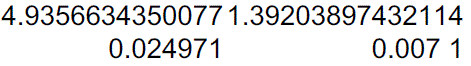

### Rate limiting steps and mechanism for the *Ndas*CDO-catalyzed reaction

*Solvent kinetic isotope effects (SKIEs) and proton inventories* – To probe the rate limiting nature of proton transfer steps, we determined SKIEs with cFG as a substrate. Fit to Equation 4 yielded ^D2O^*V*/*K*_M-cFG_ = 1.4 ± 0.2, and ^D2O^*V* = 2.1 ± 0.1. Proton inventories were complex and the best fitted model accounts for one transition state proton and one reactant state proton contributing to the observed ^D2O^*V* = 2.1 ± 0.1 (Figure S14). Due to complex nature of the enzyme filament, the fact that the fitted fractionation factors assume the ^D2O^*V* is an intrinsic isotope effect (when there is no evidence supporting or discarding this assumption), we do not discuss this further in terms of mechanism, and future studies focused on the catalytic and chemical mechanism of this reaction are needed to establish the nature of this complex effect. To rule out differences in viscosity between H_2_O and D_2_O giving rise to the observed SKIEs, we performed control experiments with 9% glycerol, as this yields a relative viscosity akin to D_2_O. No viscosity effect at this concentration was observed (Figure 5a).

**Figure 5:**
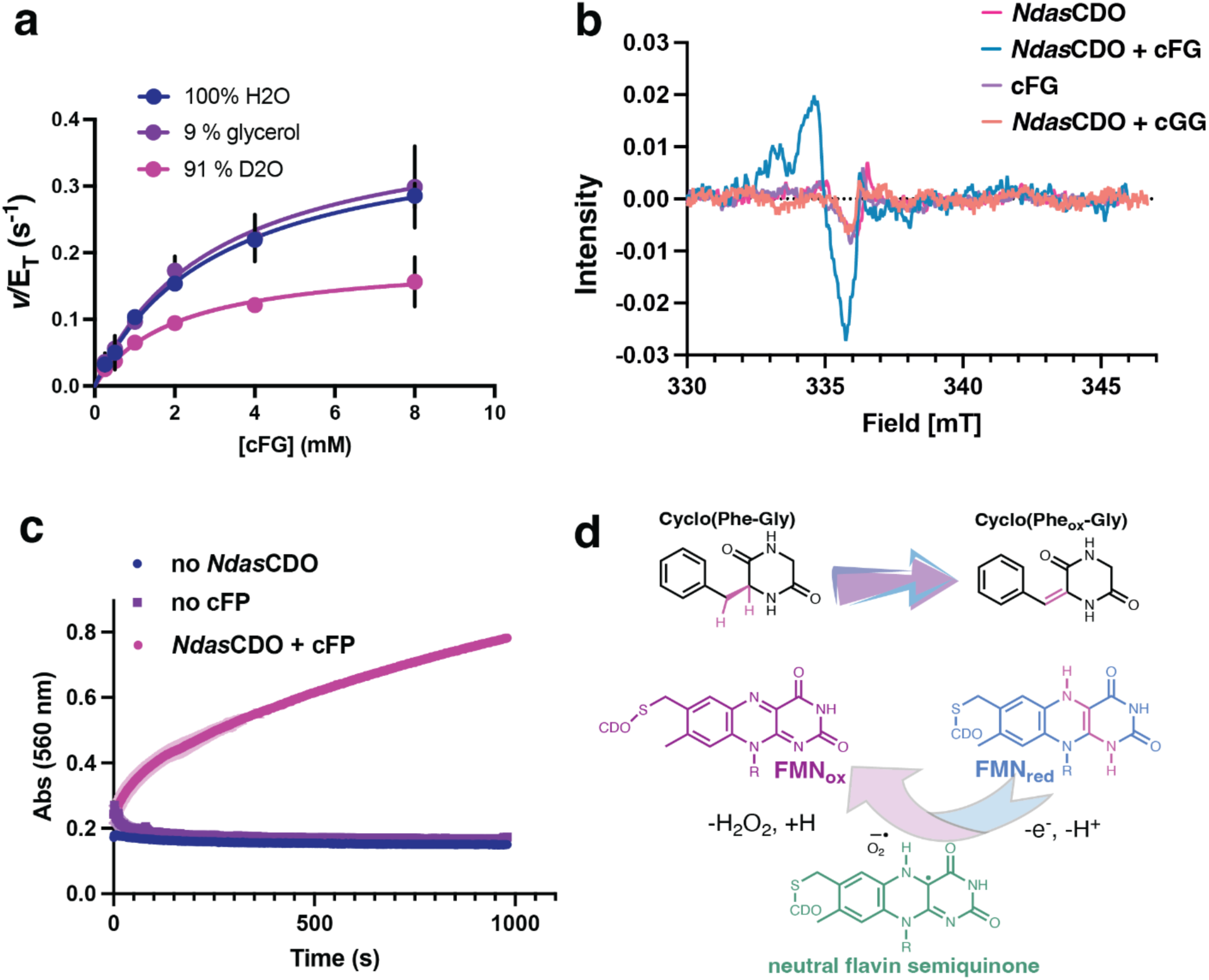
Rate limiting protonation steps and radical formation on the *Ndas*CDO-catalyzed reaction. a) Solvent kinetic isotope effects with 91 % D2O and with 9 % glycerol as a viscosity control yield ^D2O^*V*/*K*_M-cFG_ = 1.4 ± 0.2, and ^D2O^*V* = 2.1 ± 0.1. Line is a fit to Equation 4. b) Continuous wave EPR spectra acquired in samples frozen in liquid nitrogen with 5 mM cFG (blue) or cGG (orange) and 100 μM *Ndas*CDO. Controls with cFP only (purple) or *Ndas*CDO only (pink) are also shown. c) Coupled reaction with hydrogen peroxidase and ABTS (2,2’-Azinobis [3-ethylbenzothiazoline-6-sulfonic acid]-diammonium salt), showing hydrogen peroxide formation. d) Summarized catalytic cycle depicting a potential neutral flavin semiquinone intermediate.

**Figure 6:**
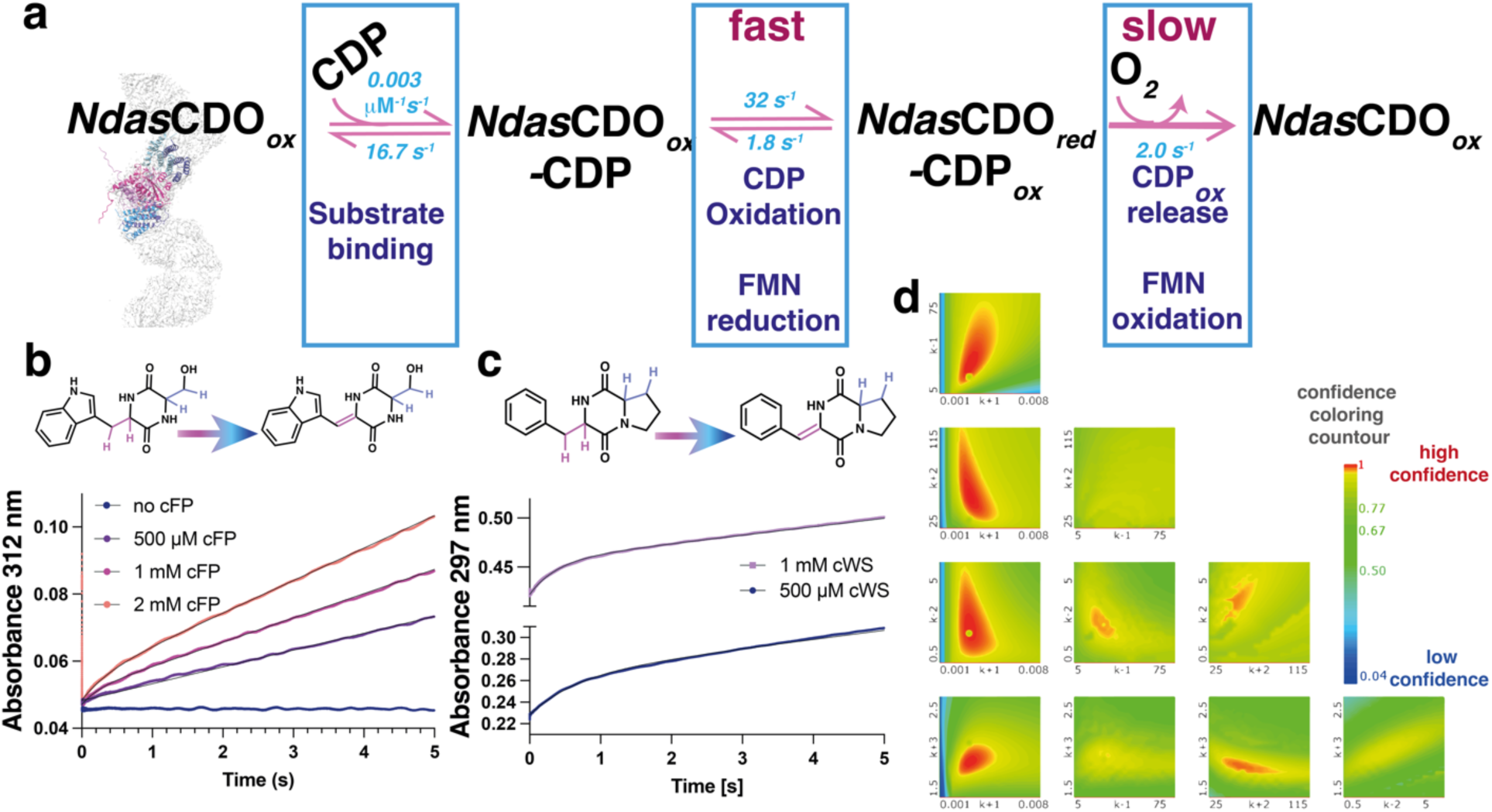
Catalytic cycle for the *Ndas*CDO-catalysed reaction. a) Model used to fit multiple turnover experiment with substrate cFP, displaying best fitted values. Boxes depict macroscopic steps, as the signal observed originated from the formation of cFP-2H. The last step in the model includes product release and FMN re-oxidation so next catalytic cycle can occur. b) Raw data for multiple turnover experiment with 8.5 μM *Ndas*CDO and varying concentrations of cFP (0.5, 1 and 2 mM), monitored at 312 nm. c) Raw data for multiple turnover experiment with 8.5 μM *Ndas*CDO and varying concentrations of cWS (0.5, and 1 mM), monitored at 297 nm. For b and c, in the time frame of the experiment we expect a single oxidation to take place, data are depicted as the average of three individual replicates and the black line is a fit to Equation 7. d) Results from data fitting using Kintek Global Explorer and output from the three-dimensional error analysis performed using Fitspace. High confidence values for the fits are depicted in red, bright yellow boundaries represent the limits of the confidence interval for each rate constant when varied in function of every other rate constant in the mechanism proposed. Table 2 summarizes the values obtained after fitting.

*Viscosity studies* – Glycerol, sucrose and PEG were used as viscogens to determine whether diffusional steps are contributing to *k*_cat_/*K*_M-cFP_ and *k*_cat_. Glycerol and sucrose are acting as microviscogens, while PEG 8K acts as a macroviscogen.^45^ No effect was observed with increasing concentrations of neither viscogen (Figure S15, slopes equal to 0.1 ± 0.1). We conclude diffusional steps included in substrate binding and product release as well as large conformational changes the filament enzyme might be undergoing during catalysis do not significantly limit catalytic turnover.^45^

*Radical formation* – as many flavin dependent enzymes have radical reaction intermediates, we used EPR to test whether this was the case for *Ndas*CDO. We observed a radical formed solely in the presence of enzyme and the cFG substrate, while no radical formation was seen in the enzyme alone or with cFG under identical experimental conditions (Figure 5b). We also did not observe radical formation with *Ndas*CDO and cyclo-Glycine-Glycine (cGG), and therefore conclude the radical observed is exclusively formed as reaction takes place. Figure 5d depicts a neutral flavin semiquinone intermediate, which is agreement with our data, however, the precise nature of this radical remains to be determined by additional work.

*Hydrogen peroxide generation* – In the CDP from *Streptomyces albulus,* hydrogen peroxide is formed during turnover.^20^ To confirm H_2_O_2_ formation in the *Ndas*CDO-catalyzed reaction we used a used a coupled assay with hydrogen peroxidase and ABTS (2,2’-Azinobis [3-ethylbenzothiazoline-6-sulfonic acid]-diammonium salt). An increase in absorbance at 560nm only in the presence of NdasCDO and cFP, consistent with H_2_O_2_ formation a reaction product was observed (Figure 5c).

*Burst of product formation* – to determine whether the first turnover of the reaction is slower than subsequent turnovers (burst of product formation), indicative of a step after the chemical step under evaluation being rate limiting, we carried out multiple turnover experiments under pre-steady state conditions. A clear burst was observed for two distinct substrates, with similar observed rate constants for the exponential phase when fitted analytically (at 1 mM both substrates *k*_obs-cWS_ = 2.6 ± 0.1 s^−1^, *k*_obs-cWS_ = 4.2 ± 0.1 s^−1^). Care must be taken comparing both substrates, as they likely have different affinities for the enzyme, but nevertheless rates for the exponential burst phase with both substrates are in the same order of magnitude.

Data acquired with cWS possesses higher signal amplitude, and therefore burst is more pronounced, but absorbance at 297 nm is complex given contributions from the FMN cofactor, the enzyme and the substrate. Alternatively, cFP dehydrogenation leads to increase in absorbance at 312 nm, which is unique given cFP has no absorbance at this wavelength, the contribution from the enzyme is less significant, enabling data analysis using a more complex model and numerical fitting. Our three step model accounts for substrate binding, CDP oxidation and FMN oxidation/product release. Because we don’t have observable signals to discriminate individual rate constants for FMN oxidation and product release in individual steps, they were combined as a single irreversible apparent rate constant. However, the absence of viscosity effects and observation of a sizeable solvent kinetic isotope effect on *k*_cat_ point towards a step coupled to FMN re-oxidation as rate limiting, instead of product diffusion from enzyme. This can be directly linked to FMN re-oxidation or be due to a proton-transfer-linked conformational change, as proposed for the phosphoribosyl-ATP Pyrophosphohydrolase/Phosphoribosyl-AMP Cyclohydrolase (AbHisIE) from *Acinetobacter baumanii*^46^.

## Conclusions

*Biocatalysis using CDO enzymes –* Oxidized cyclic peptide natural products such as phenylahistin^6^ and albonoursin^21^ have anticancer activity, and the importance of dehydroamino acid residues in bioactive natural products has been established.^47^ Therefore, CDO enzymes can be useful biocatalysts, provided they are better understood in terms of substrate selection and properties.

Performing dehydrogenation en masse by utilising a CDO biocatalyst would maximise atom economy whilst minimising unwanted side products and waste from this selective reaction. Previous attempts at mutating enzymes that catalyse reactions via radical intermediates successfully expanded the overall substrate scope^48^ for improved biocatalysts. Furthermore, previous work by our group^44^ and others^49^ established that the biosynthetic partner of CDOs, cyclodipeptide synthases, were amenable to active site manipulation for expanded substrate acceptance^44^, and that these enzymes were capable of utilizing unnatural amino acids^50, 51^. Other tailoring enzymes have been described, which catalyze prenylation^52^ as well as other complex reactions^53^. Thus, our work in better understanding the structure and kinetics of *Ndas*CDO is a necessary step towards combining these strategies to generate novel cyclodipeptide analogues.

*Mechanism, structure and catalytic turnover by a filament enzyme –* Here we report details on the mechanism and structure of a cryptic member of the cyclodipeptide oxidase family. In agreement with recent findings by *Giessen et al.*^41^, *Ndas*CDO was observed as a high molecular weight oligomer whose length is dependent on overall salt concentration. *Ndas*CDO is an enzyme from the halophile *Nocardiopsis dassonvillei*, and therefore the environment naturally occupied by *N. dassonvillei* could be conducive to filament formation and generation of oxidized CDPs. Prior work with ethyl acetate extracts of this actinobacteria demonstrated biofilm-inhibitory activity, although the precise identity of the natural product(s) giving rise to this effect remains to be determined^54^. Previous research discussed the inactivation of CDOs when either subunit was absent, and we demonstrate structural features underlying this reliance on a functional complex. Filament formation is fundamental for enzymatic activity, as our structural work shows residues from subunits A and B are important to interact with substrates and the covalently bound FMN cofactor.

Whilst *Ndas*CDO-A harbours high structural similarity to nitroreductases, our work highlights that the active site of CDOs differs from these enzymes, as it does not rely on the same catalytic mechanism and structural features. Filamentation could aid *Ndas*CDO in evading degradation during starvation or stress^55^ – both environmental conditions that *Nocardiopsis dassonvillei* thrives in^54^. Filament enzymes are emerging as an important concept in the regulation of protein function^56^, and therefore our work sheds light into how structure and function are amalgamated to control activity in one of these complex systems.

## Supporting Information

The Supporting Information is available free of charge and includes additional methods, supporting Figures and Tables. Table S3 contains information about the cryo-EM structure deposited here.

## Accession Codes

*Ndas*1146 (Uniprot: D7B1W6) and *Ndas*1147 (Uniprot: D7B1W7), pdb accession code 9EXV and EMDB accession code EMD-50049.

## Data availability statement

Data for kinetic assays is available in the supporting information file, as well as NMR spectra and spectra for determination of extinction coefficients used. Raw mass spectrometry data is available on the Figshare project “<u>Broad substrate scope C-Coxidation in cyclodipeptides catalysed by a flavin-dependent filament</u>” https://doi.org/10.6084/m9.figshare.25556850

## AUTHOR INFORMATION

### Author Contributions

Manuscript was written by first author in its majority, but all authors contributed to the final form. All authors have given approval to the final version of the manuscript. Specific contributions are as follows:

**Emmajay Sutherland**: Designed and performed experiments, interpreted data, wrote manuscript.

**Christopher Harding**: contributed to protein crystallography experiments, interpreted data, revised manuscript.

**Tancrede Martin Y Du Monceau De Bergendal** (T.M.M.B) synthesized cWS and contributed to activity assays, analyzed data, revised manuscript.

**Gordon J. Florence** supervised T.M.M.B, analyzed data, revised manuscript.

**Katrin Ackermann**: performed EPR experiments, analyzed data, revised manuscript.

**Silvia Sinowski**: performed protein mass spectrometry experiments, analyzed data, revised manuscript.

**Bela Bode**: designed and performed EPR experiments, analyzed data, revised manuscript.

**Ramasubramanian Sundaramoorthy**: Prepared cryo-EM grids, acquired data, analyzed data, revised manuscript.

**Clarissa Melo Czekster:** participated in project conception, analyzed and interpreted data, revised manuscript.

## ACKNOWLEDGMENT

We acknowledge the University of Dundee Cryo-EM facility for access to the instrumentation, funded by Wellcome (223816/Z/21/Z) and MRC (MRC World Class Laboratories PO 4050845509). We thank electron Bio-Imaging Centre (eBIC) facility, Diamond light source Ltd, UK for collection of 300kV electron microscope data.

## ABBREVIATIONS

EPR: Electron paramagnetic resonance
LC-MS: liquid chromatography mass spectrometry
*Ndas*CDO: cyclodipeptide oxidase from *Nocardiopsis dassonvileii*
CDPS: cyclodipeptide synthase
CDP: cyclodipeptide. All cyclic dipeptides used here were composed by L-amino acids and abbreviated with the letter “c” for cyclo, followed by the 1 letter code for each amino acid as follows: cFG, cFP, cFF, cLP, cHF, cWG, cWW, cWY, cWS

