## Supporting information for "Broad substrate scope C-C oxidation in cyclodipeptides catalysed by a flavin-dependent filament"

***To whom correspondence should be addressed:**

### Additional methods

#### *Ndas*CDO plasmid map

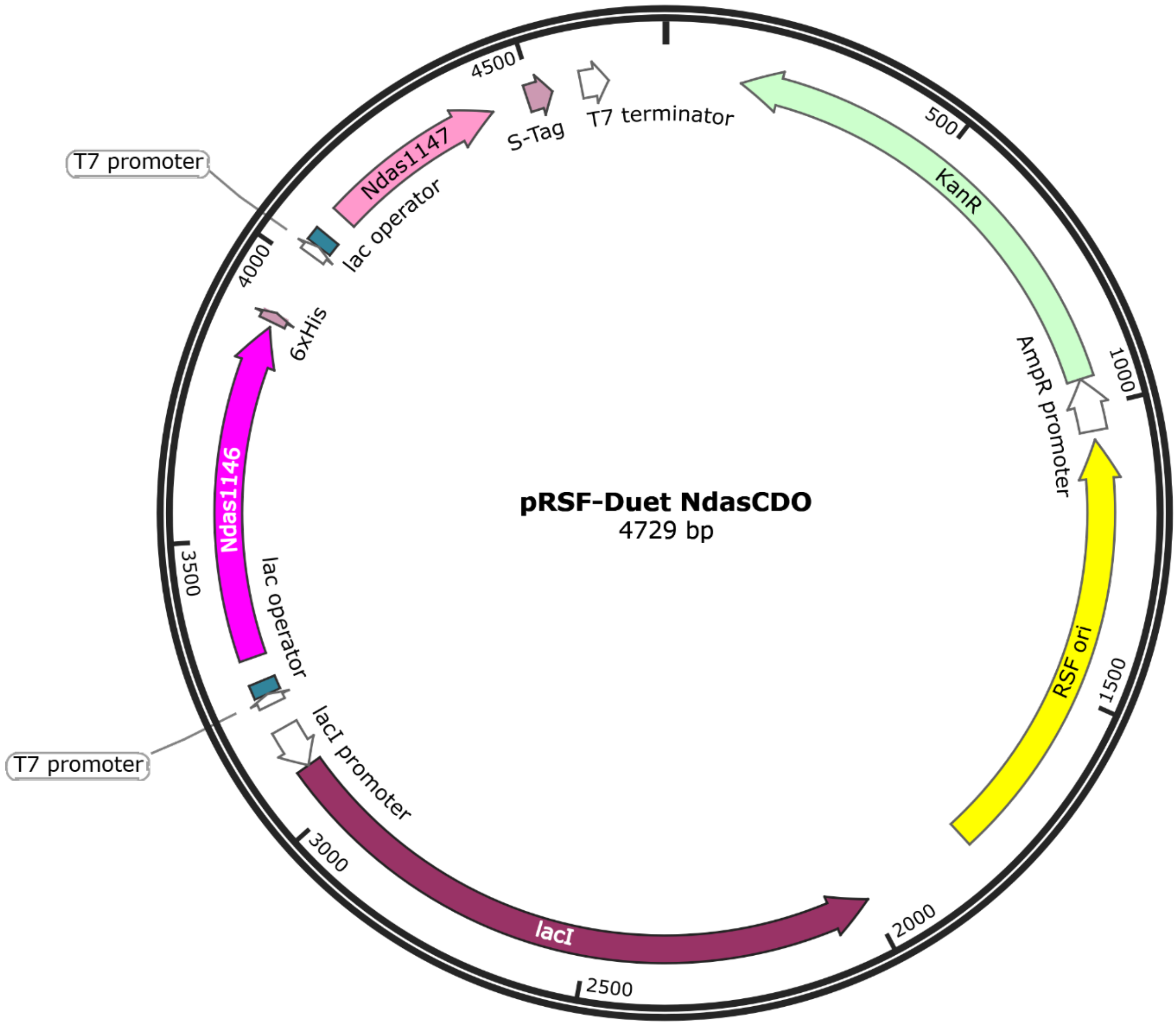

#### *Ndas*CDO and mutant-*Ndas*CDO protein sequences

*Ndas*1146

MDTGSSEPDANRCPSQRSSHALQTLTTRRAVRAFADRPVDDSLLDPMLDAMLAAP
SASNKQAWAFVAVRERRALRLLRAFSPGIIELPPLVVAACFDRSRAVGGSGNSTDS
GDSWDEGMLCVAMAVENLLLAAHCLGLGGCPSGSFRRGPVRRLLGLPDHLEPLLL
VPIGHPARPLAPAPRRDRNEVVSHERWGT

*Ndas*1147

MSAGEPEVRQVGEELLLLAAYLLSSGRGLLDEPRQYGTFRCLDAARRVLALAAGT
GPHHPELDALRGRMDDVMCGPMGDHELDTLLDQMCERLATVLEDPDVISD

*Ndas*1146-S58A

MDTGSSEPDANRCPSQRSSHALQTLTTRRAVRAFADRPVDDSLLDPMLDAMLAAP
SAANKQAWAFVAVRERRALRLLRAFSPGIIELPPLVVAACFDRSRAVGGSGNSTDS

GDSWDEGMLCVAMAVENLLLAAHCLGLGGCPSGSFRRGPVRRLLGLPDHLEPLLL VPIGHPARPLAPAPRRDRNEVVSHERWGT

*Ndas*1146 2-17 truncation

MSSHALQTLTTRRAVRAFADRPVDDSLLDPMLDAMLAAPSASNKQAWAFVAVRER RALRLLRAFSPGIIELPPLVVAACFDRSRAVGGSGNSTDSGDSWDEGMLCVAMAV ENLLLAAHCLGLGGCPSGSFRRGPVRRLLGLPDHLEPLLLVPIGHPARPLAPAPRR DRNEVVSHERWGT

*Ndas*1146 2-39 truncation

MDDSLLDPMLDAMLAAPSASNKQAWAFVAVRERRALRLLRAFSPGIIELPPLVVAA CFDRSRAVGGSGNSTDSGDSWDEGMLCVAMAVENLLLAAHCLGLGGCPSGSFR RGPVRRLLGLPDHLEPLLLVPIGHPARPLAPAPRRDRNEVVSHERWGT

#### Assay conditions

– Prior work indicated a direct absorbance change occurs upon cyclic dipeptide oxidation. We expanded this direct assay with multiple CDP substrates, since the nature of amino acid side chains changes the maximum absorbance observed from substrates and products. Using the difference spectra between reactants and products, we obtained a DeltaAbs value, which was employed to convert changes in absorbance into changes in concentration as a function of time. Substrates containing Phe, Tyr and Trp had significantly red-shifted spectra in comparison to substrates containing only non-aromatic groups (Leu, Pro, His). Figure S2 summarizes all substrates tested, with their absorption spectra, and, following overnight reactions, spectra for oxidized products (identity verified by LC-MS). A complicating factor for this assay is the fact that some CDPs can undergo two oxidation events, which can alter both kinetics and spectroscopic characteristics of intermediates. Under steady-state conditions, with excess unoxidized substrates, we observed the accumulation of a singly-oxidized product before formation of a doubly oxidized CDP in the case of cFP (Figure S8), and therefore conclude *Ndas*CDO operates distributively, catalyzing one oxidation event to generate an intermediate, which was released and could subsequently re-bind to the enzyme for a second oxidation event.

Data analysis for pH-rate profiles – According to Cook and Cleland (page 335), when two p$K_a$ values are close together, a different equation might be required to fit data, as below:

$$y = log\left(\frac{C}{1+\frac{pKa2}{pKa1}+\frac{10^{-pH}}{10^{-pKa1}}+\frac{10^{-pka2}}{10^{-pH}}}\right) \qquad \text{Equation S1}$$

However, if $K_2/K_1 < 0.1$, Equation 3 can be used, as the p$K_a$ values are distant enough to allow individual fitting. In the case of *Ndas*CDO, K2/K1 = 0.03, and the pKa values can be individually fitted.

### Supporting Figures

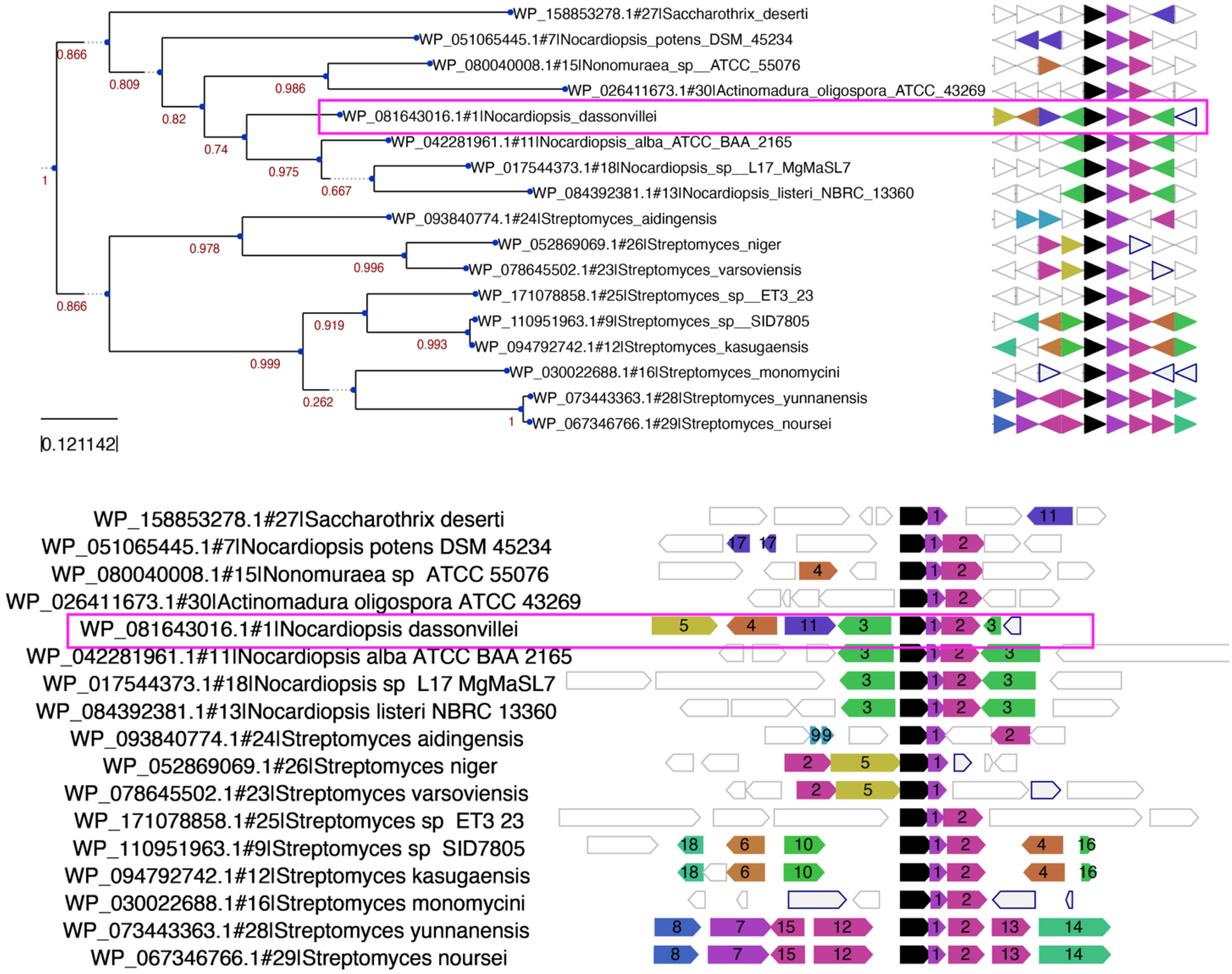

| # | Example protein | Predicted function |
|---|---|---|
| 1 | WP_238413131.1 | DUF6092 family protein (CDO subunit B) |
| 2 | WP_157851683.1 | tRNA-dependent cyclodipeptide synthase |
| 3 | WP_232532480.1 | methyltransferase |
| 4 | WP_094801803.1 | SDR family NAD(P)-dependent oxidoreductase |
| 5 | WP_094801801.1 | cytochrome P450 |
| 6 | WP_052853172.1 | MULTISPECIES: VWA domain-containing protein |
| 7 | WP_073443366.1 | MULTISPECIES: cysteine desulfurase family protein |
| 8 | WP_073443368.1 | thioesterase family protein |
| 9 | WP_093840776.1 | DUF397 domain-containing protein |
| 10 | WP_094792743.1 | HAD family hydrolase |

| | | |
|---|---|---|
| 11 | WP_158853282.1 | alpha/beta hydrolase |
| 12 | WP_189854609.1 | tRNA 2-thiouridine(34) synthase MnmA |
| 13 | WP_073443360.1 | VC0807 family protein |
| 14 | WP_073443358.1 | MULTISPECIES: NADP-specific glutamate dehydrogenase |
| 15 | WP_073443723.1 | MULTISPECIES: N-acetylmuramoyl-L-alanine amidase |
| 16 | WP_205380085.1 | MULTISPECIES: hypothetical protein |
| 17 | WP_017593943.1 | hypothetical protein |
| 18 | WP_094792745.1 | helix-turn-helix domain-containing protein |

Figure S1: Genomic context for NdasCDO. Generated using Webflags (https://server.atkinson-lab.com/webflags)[1]. Top, phylogenetic tree of aligned sequences. Bottom, biosynthetic gene clusters with closest homologues of *Ndas*1146 (Uniprot: D7B1W6, shown in black).

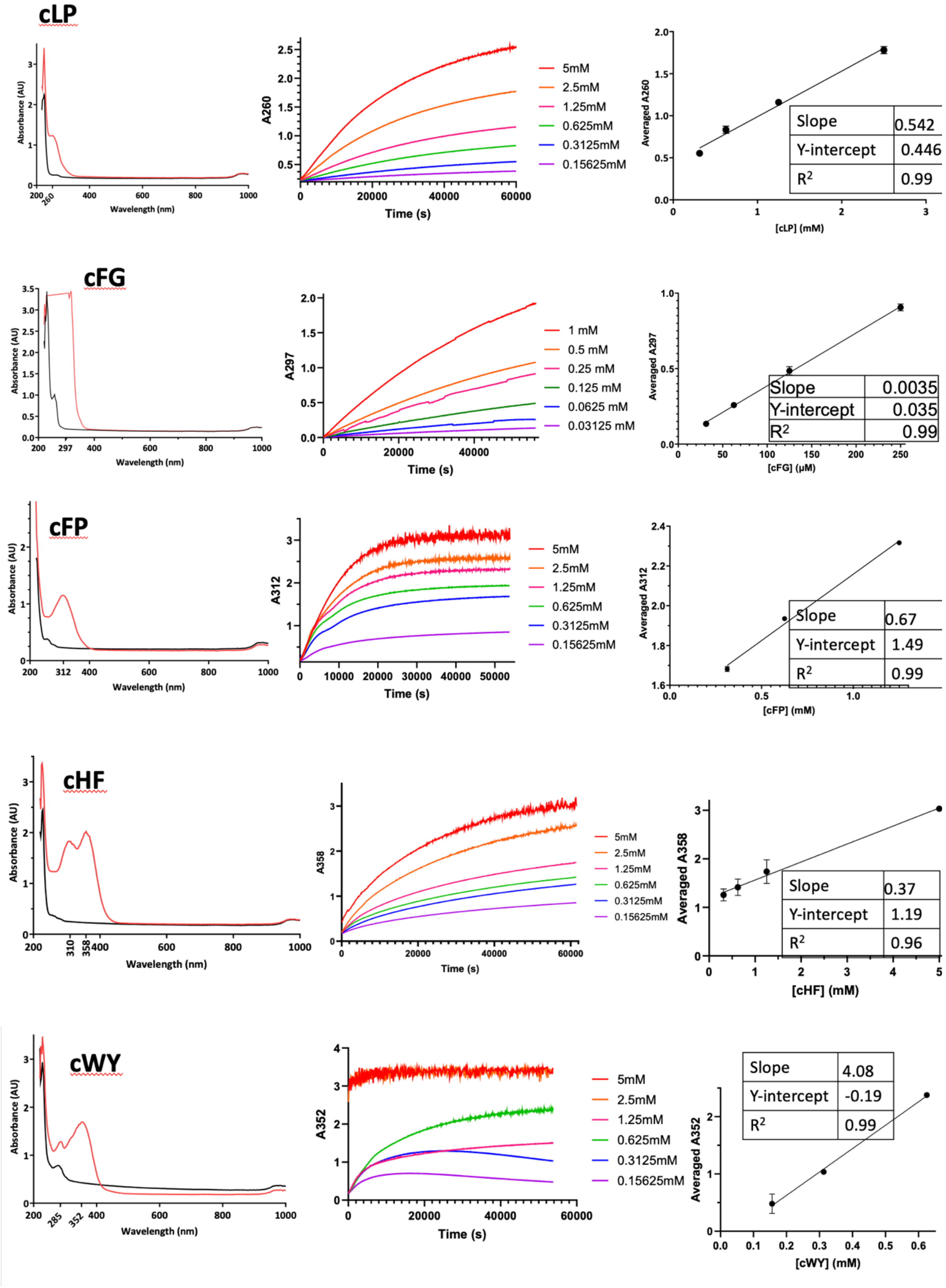

cLP
Absorbance (AU)
Wavelength (nm)
A260
Time (s)
5mM
2.5mM
1.25mM
0.625mM
0.3125mM
0.15625mM
Averaged A260
[cLP] (mM)
Slope 0.542
Y-intercept 0.446
R2 0.99
cFG
Absorbance (AU)
Wavelength (nm)
A297
Time (s)
1 mM
0.5 mM
0.25 mM
0.125 mM
0.0625 mM
0.03125 mM
Averaged A297
[cFG] (μM)
Slope 0.0035
Y-intercept 0.035
R2 0.99
cFP
Absorbance (AU)
Wavelength (nm)
A312
Time (s)
5mM
2.5mM
1.25mM
0.625mM
0.3125mM
0.15625mM
Averaged A312
[cFP] (mM)
Slope 0.67
Y-intercept 1.49
R2 0.99
cHF
Absorbance (AU)
Wavelength (nm)
A358
Time (s)
5mM
2.5mM
1.25mM
0.625mM
0.3125mM
0.15625mM
Averaged A358
[cHF] (mM)
Slope 0.37
Y-intercept 1.19
R2 0.96
cWY
Absorbance (AU)
Wavelength (nm)
A352
Time (s)
5mM
2.5mM
1.25mM
0.625mM
0.3125mM
0.15625mM
Averaged A352
[cWY] (mM)
Slope 4.08
Y-intercept -0.19
R2 0.99

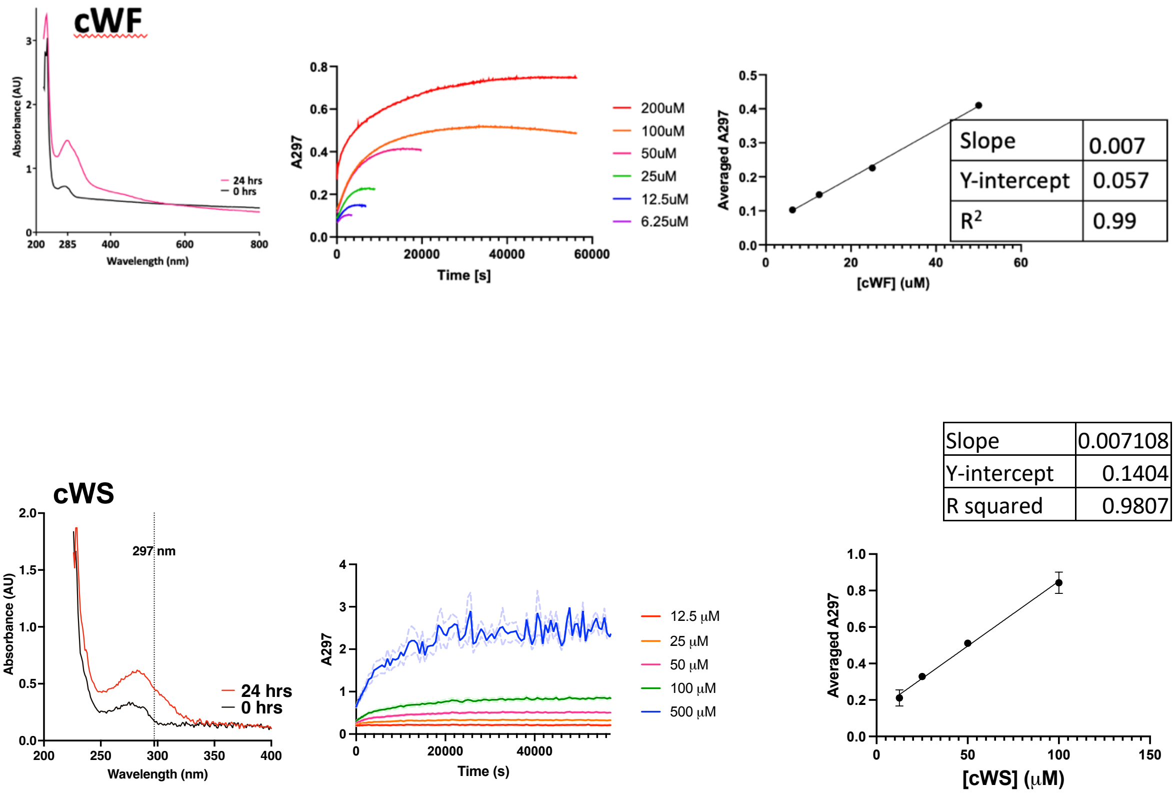

Figure S2: UV difference spectra upon CDP oxidation. Left: difference UV spectra comparing substrates (black trace) and oxidized products after reaction reached completion (red trace); middle: progress curve for each rection until completion, endpoint UV reading was used to generate the calibration curve on the right, to obtain a relationship between absorbance change and CDP oxidation for each CDP substrate.

### Figure S3: Test Reactions with distinct CDP substrates.

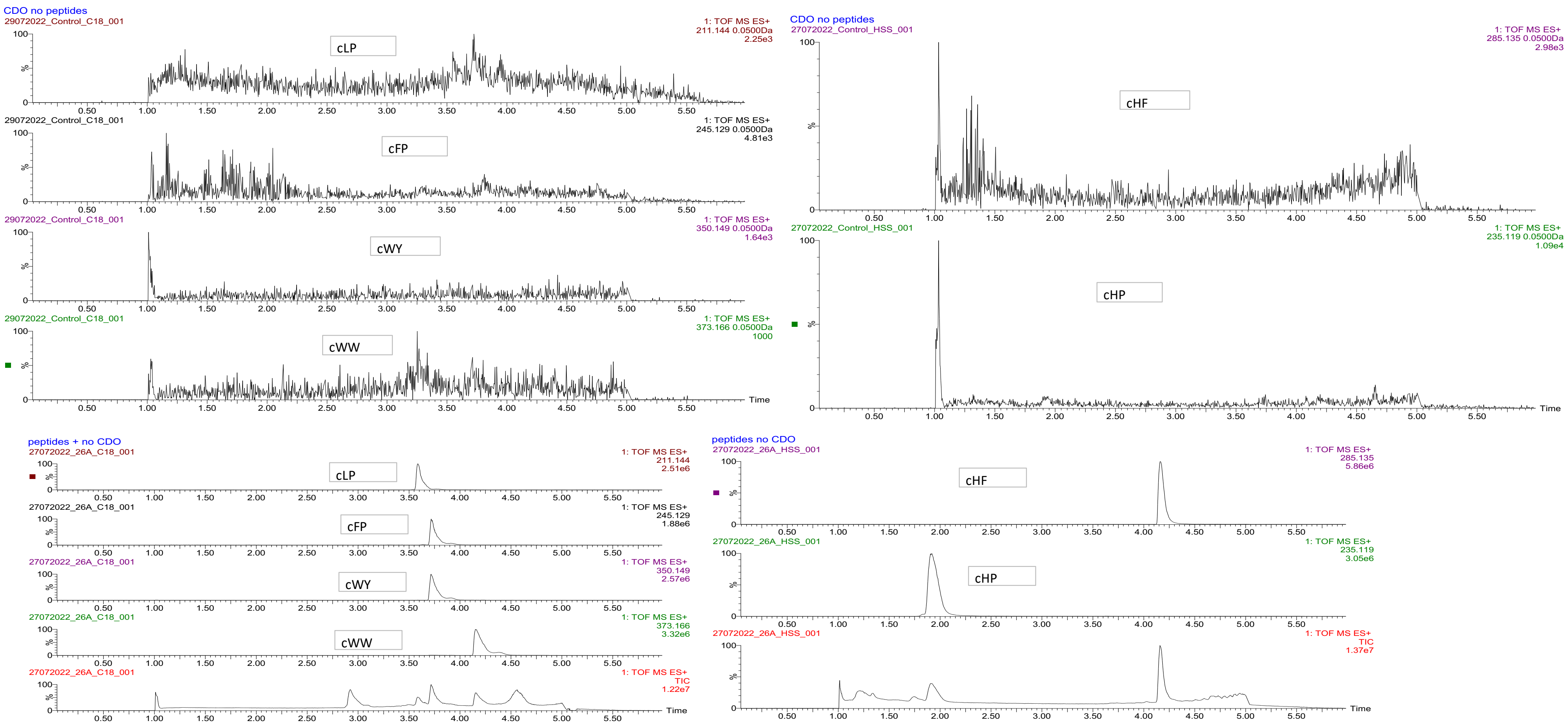

Figure S3a) Negative control reactions with Distinct CDP substrates.

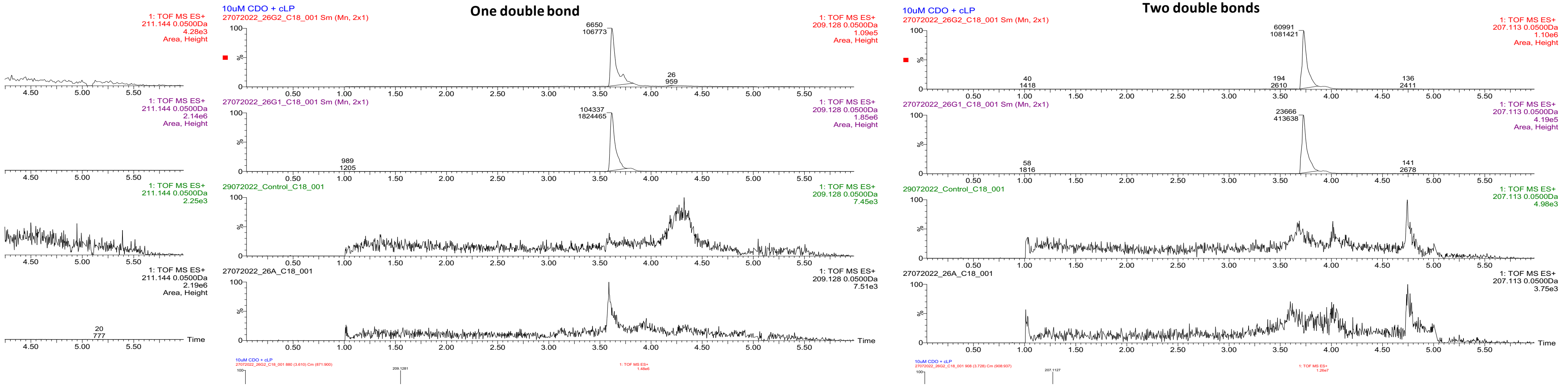

Figure S3b) Raw data for overnight reaction with cLP.

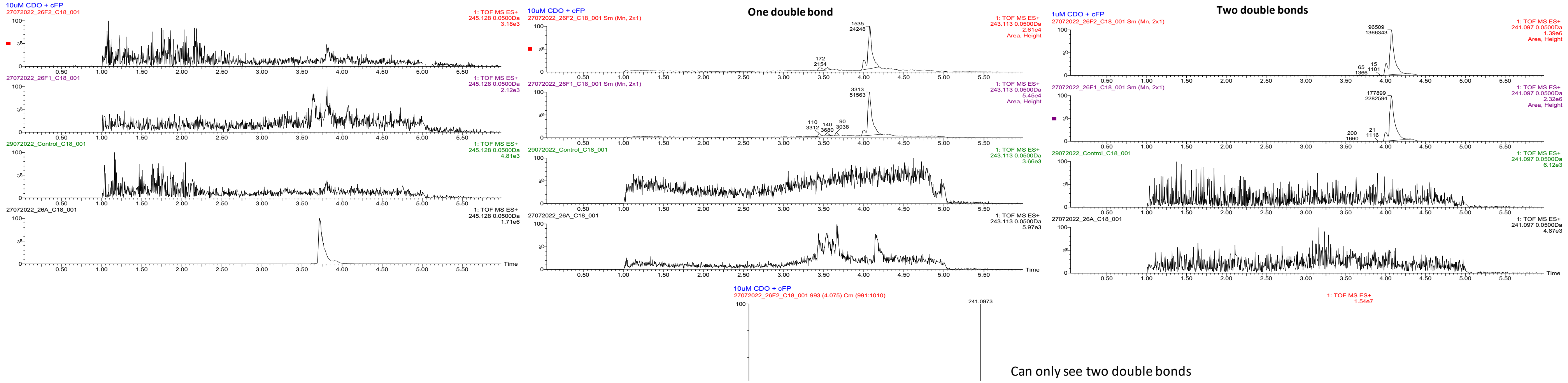

Figure S3c) Raw data for overnight reaction with cFP.

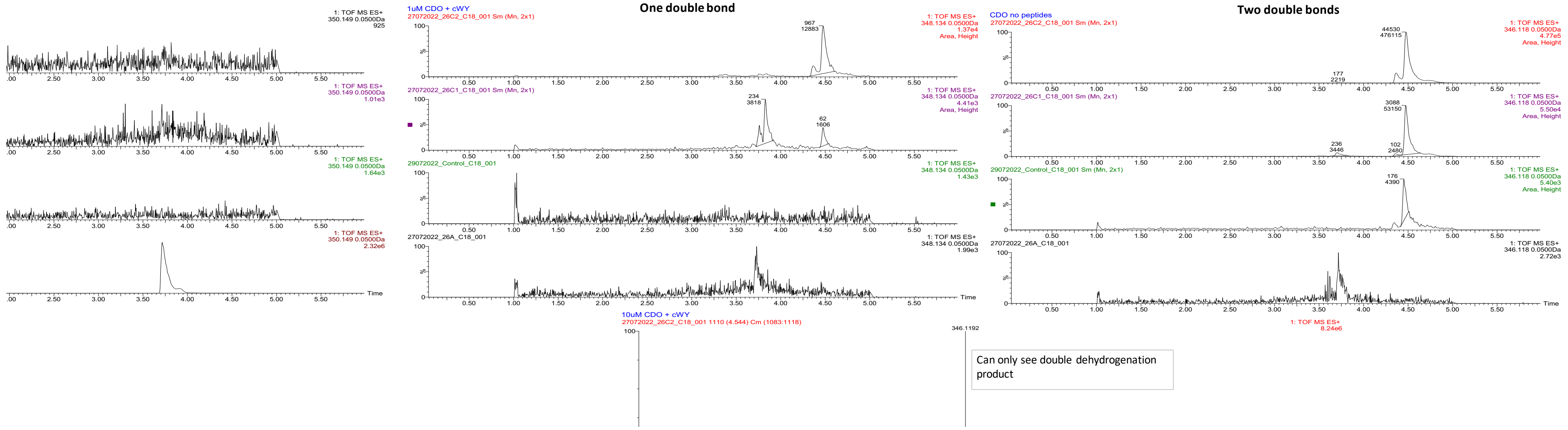

Figure S3d) Raw data for overnight reaction with cWY.

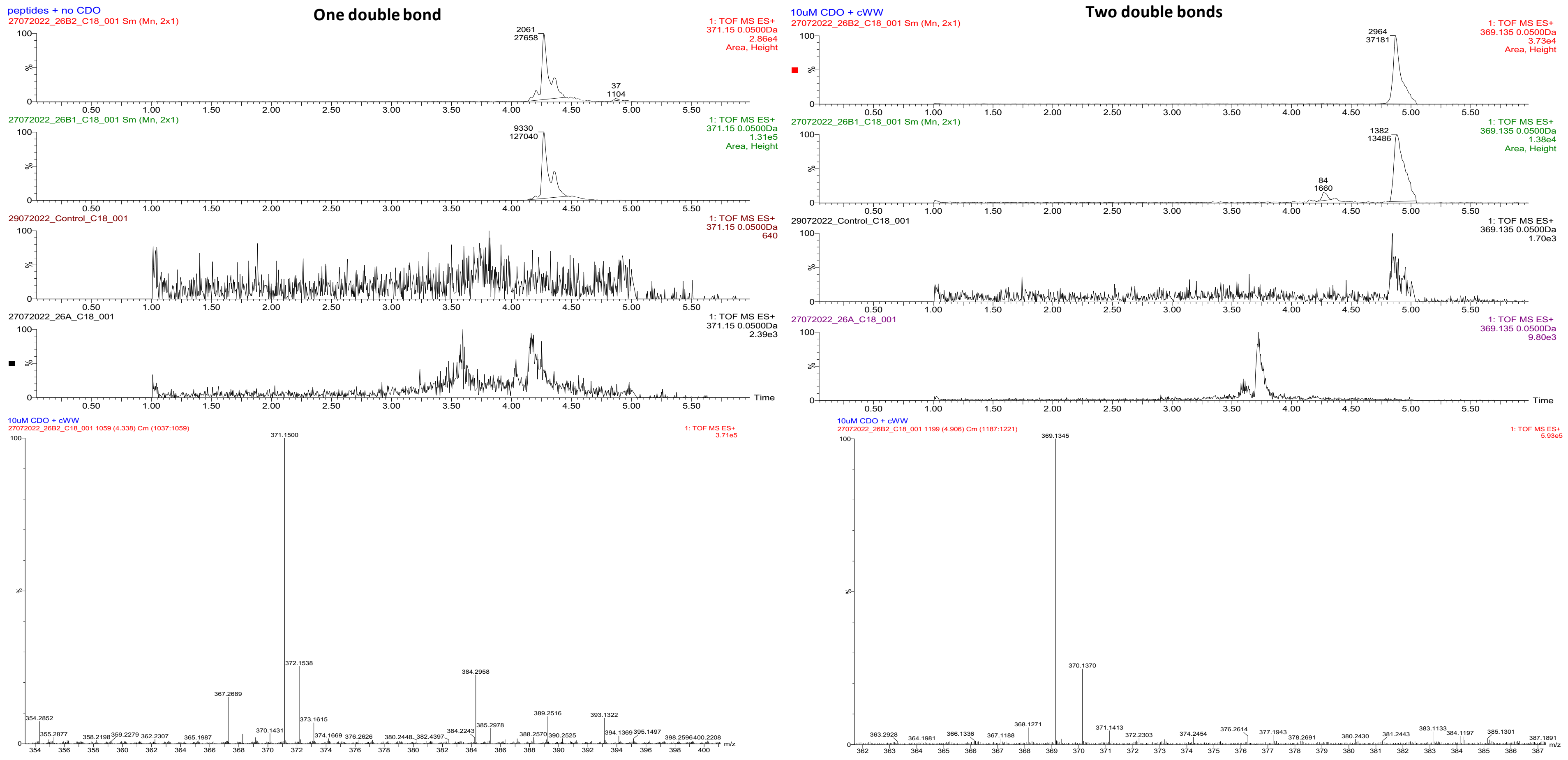

Figure S3e) Raw data for overnight reaction with cWW.

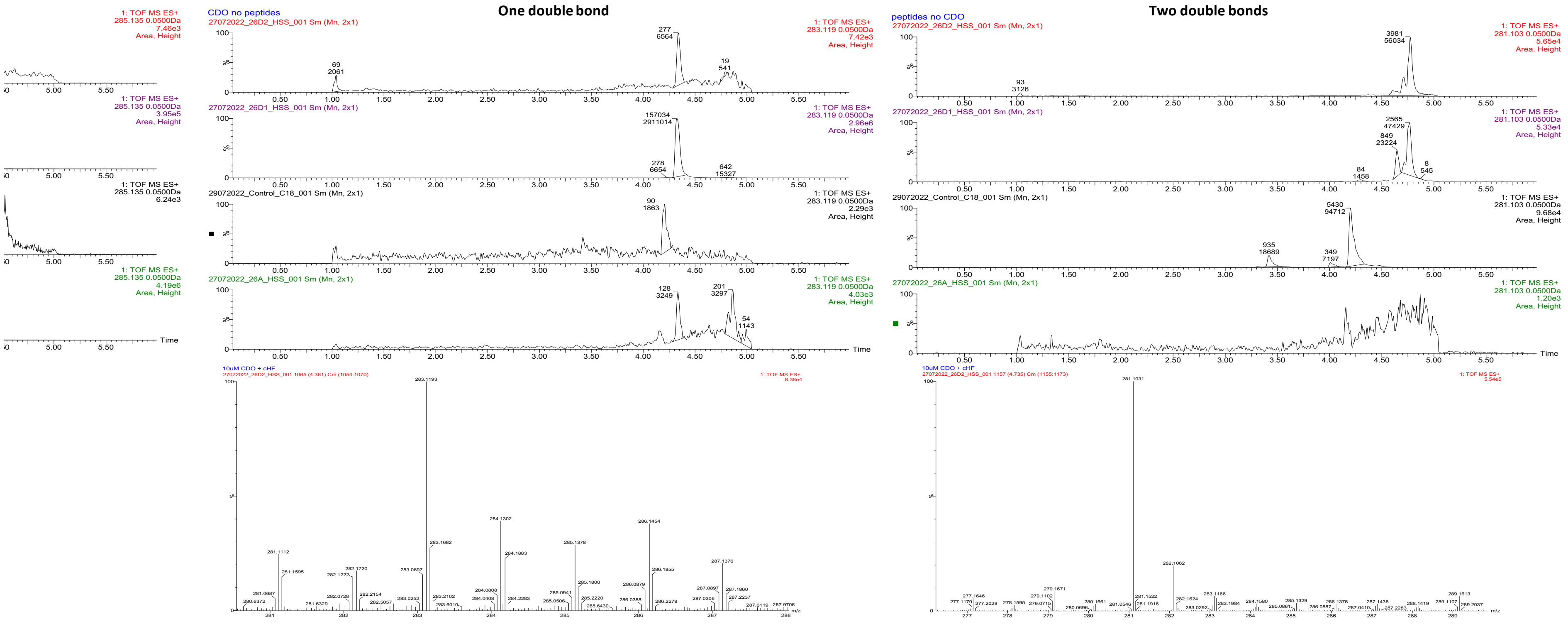

Figure S3f) Raw data for overnight reaction with cHF.

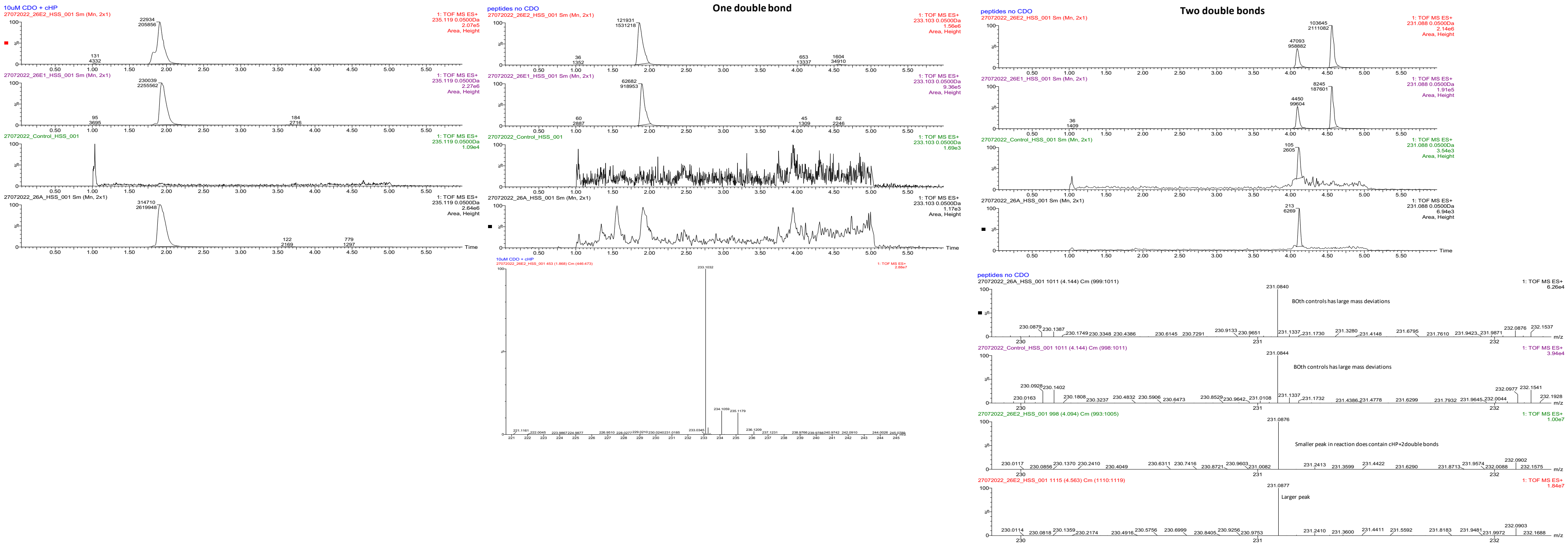

Figure S3g) Raw data for overnight reaction with cHP.

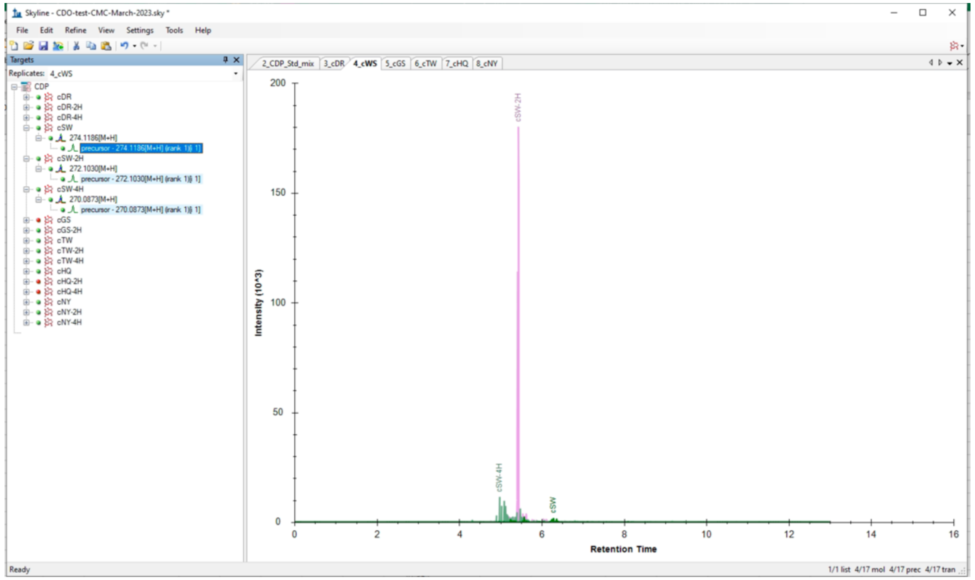

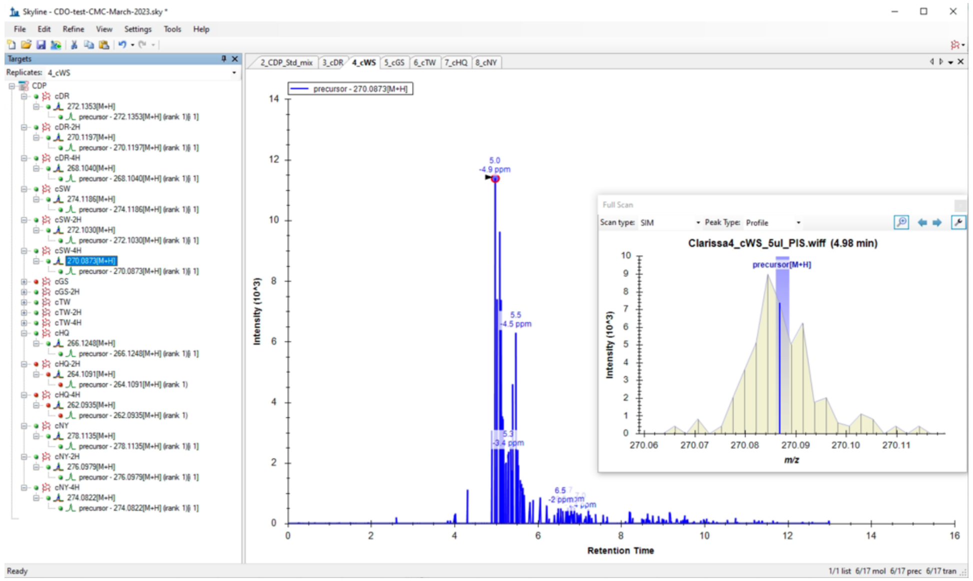

Figure S3h) Raw data for overnight reaction with cWS. Data was analysed using Skyline[2], compared to standards and confidence set to match masses with <5 ppm deviation from expected theoretical mass.

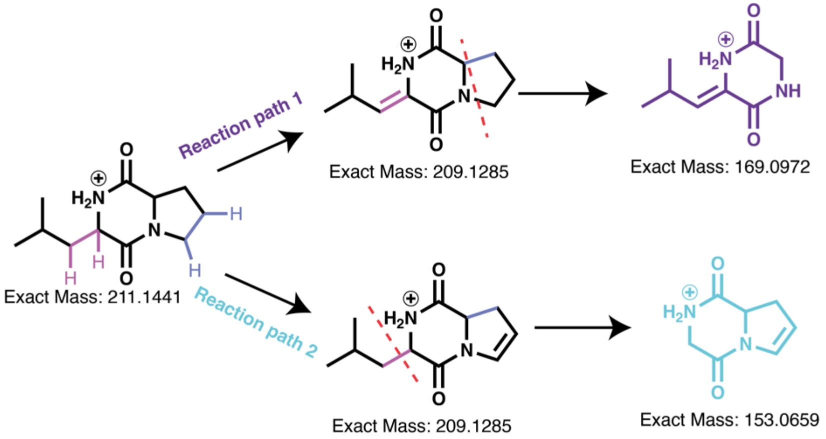

Figure S3i) Unique fragmentation products expected after fragmentation of the product cLP-2H.

A)

B)

C)

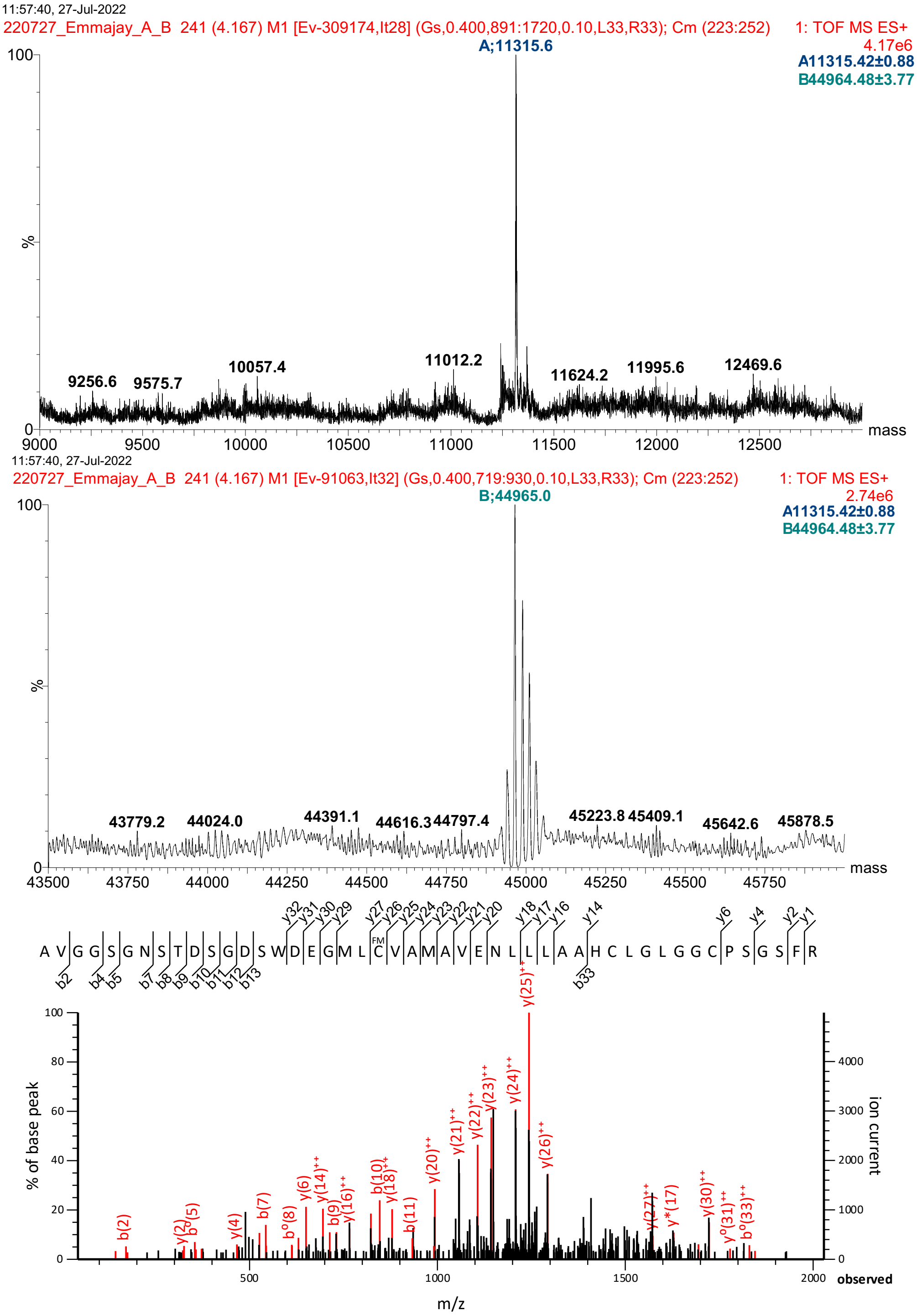

**Figure S4: Protein mass spectrometry to confirm covalently bound FMN.** Protein intact mass for subunit A) shows no modifications on subunit B. B) shows subunit A presents as a dimer and is modified by the addition of FMN. C) Peptides observed after trypsin digestion, showing FMN is covalently attached to C121 of subunit A.

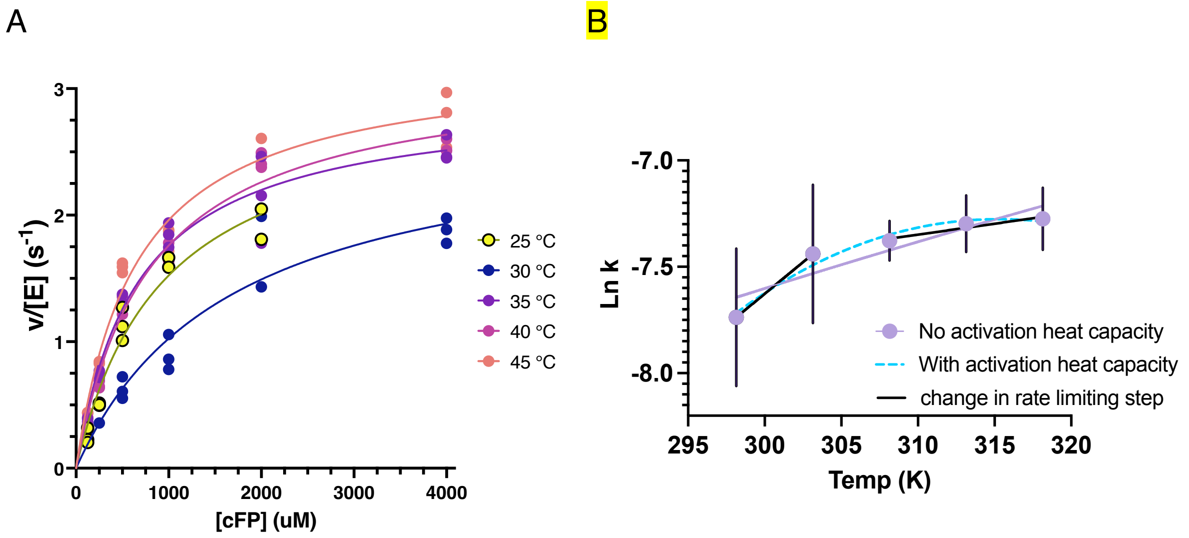

#### Figure S5: Temperature rate profiles.

A) raw data for Michaelis-Menten plots at each different temperature. All data are shown.

b) Comparison between fits to an Eyring Equation with and without contributions from activation heat capacity, as well as with linear fits that would occur due to changes in the rate limiting step for the reaction under study.

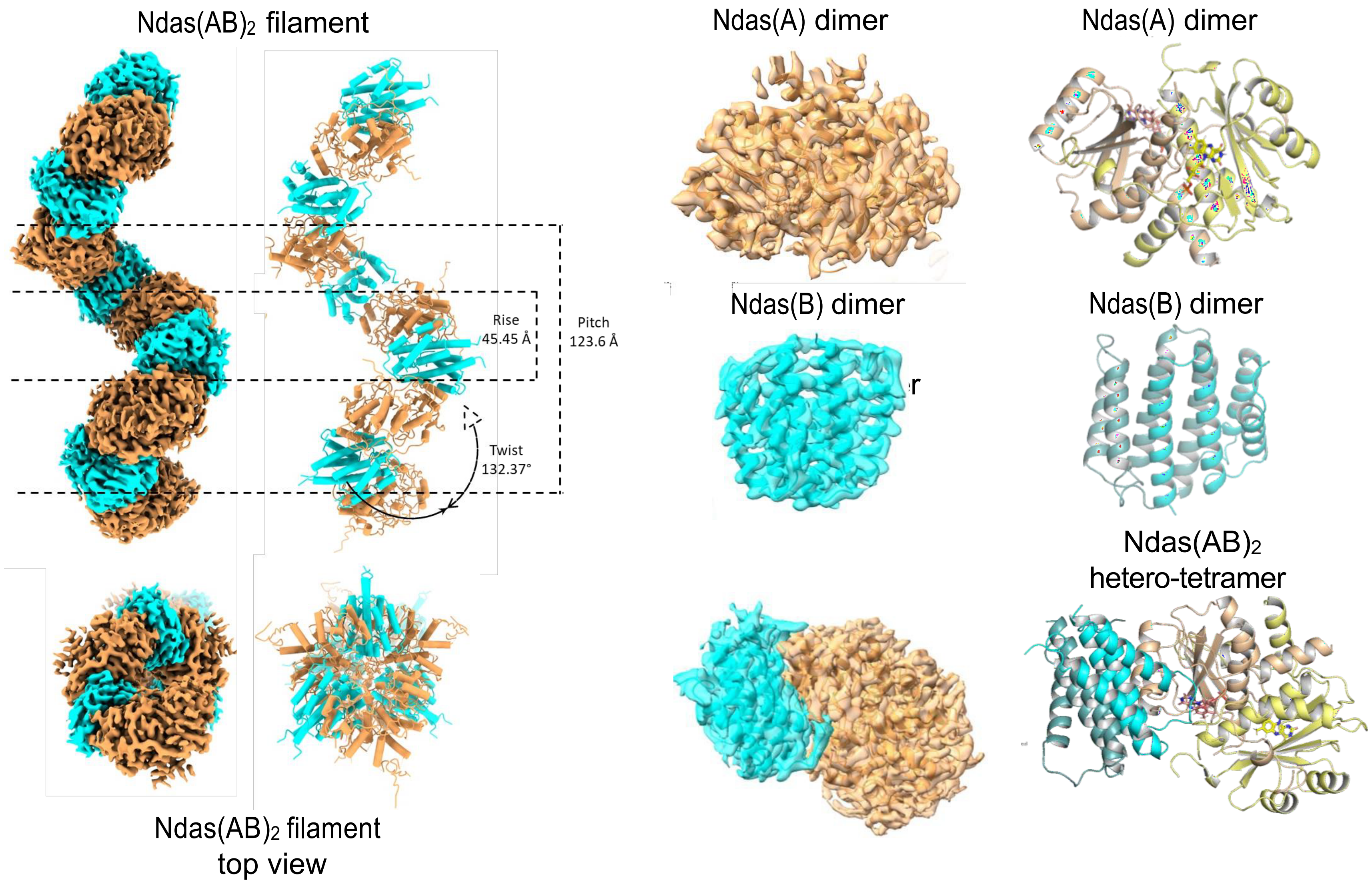

**Figure S6: Overview of cryo-EM structure.** A subunits are in tan, B subunits in cyan. a) overview of filament; b) Maps for subunits (left) as well the cartoon representation. Cryo-EM models for subunits alone and in combination (bottom depicts the dimer of dimers).

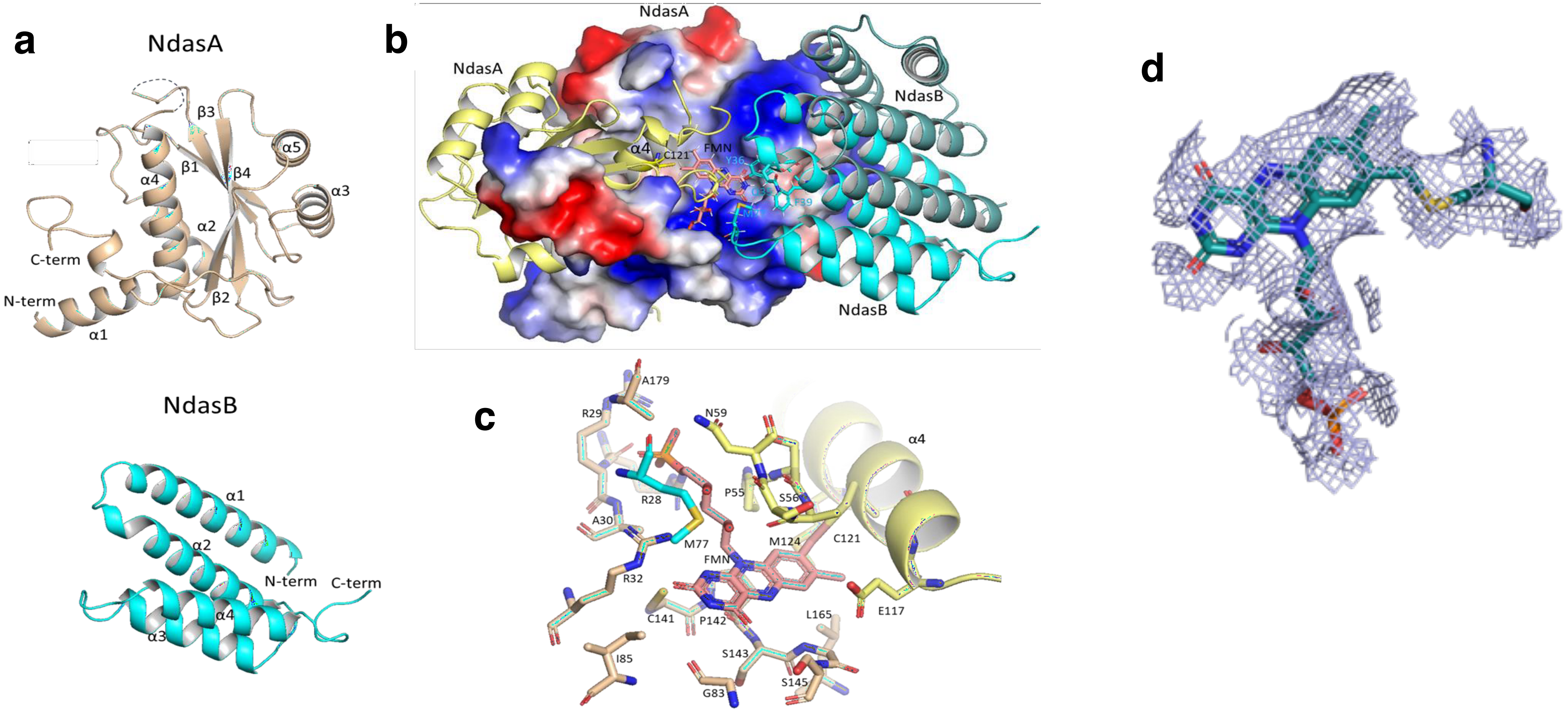

Figure S7: Details on the FMN binding site and overall structure of A and B subunits. a) Topology for individual A and B subunits. b) Electrostatic potential surface surrounding the FMN binding site. c) Residues in the vicinity of the FMN binding site. d) Density map surrounding the FMN cofactor and showing continuous density towards Cys121. Map shown at 2.0 s contour, prepared using Pymol.

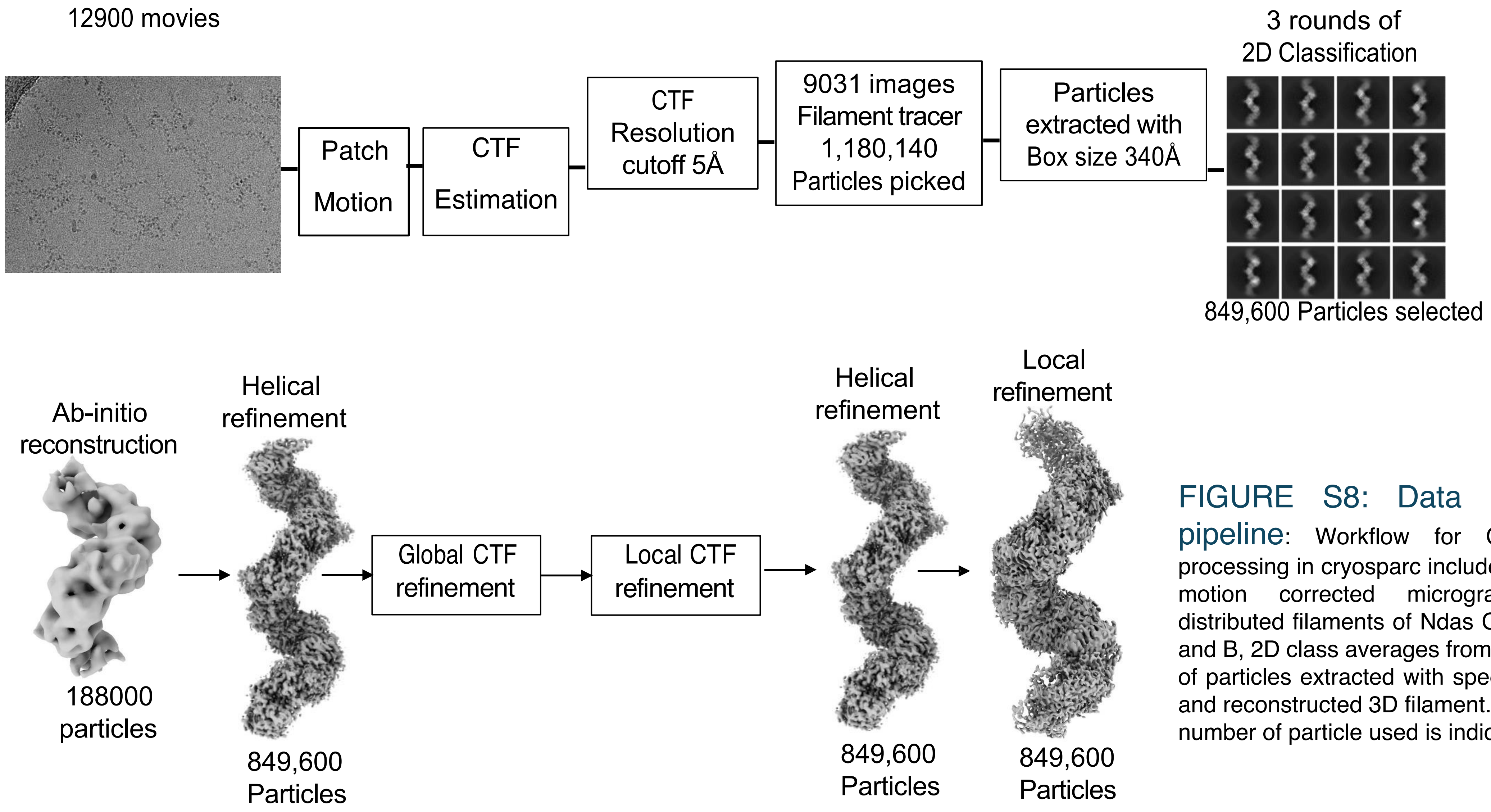

FIGURE S8: Data processing pipeline: Workflow for CryoEM data processing in cryosparc includes an example motion corrected micrograph showing distributed filaments of Ndas CDO subunit A and B, 2D class averages from cleaned stack of particles extracted with specified box size and reconstructed 3D filament. At each stage number of particle used is indicated.

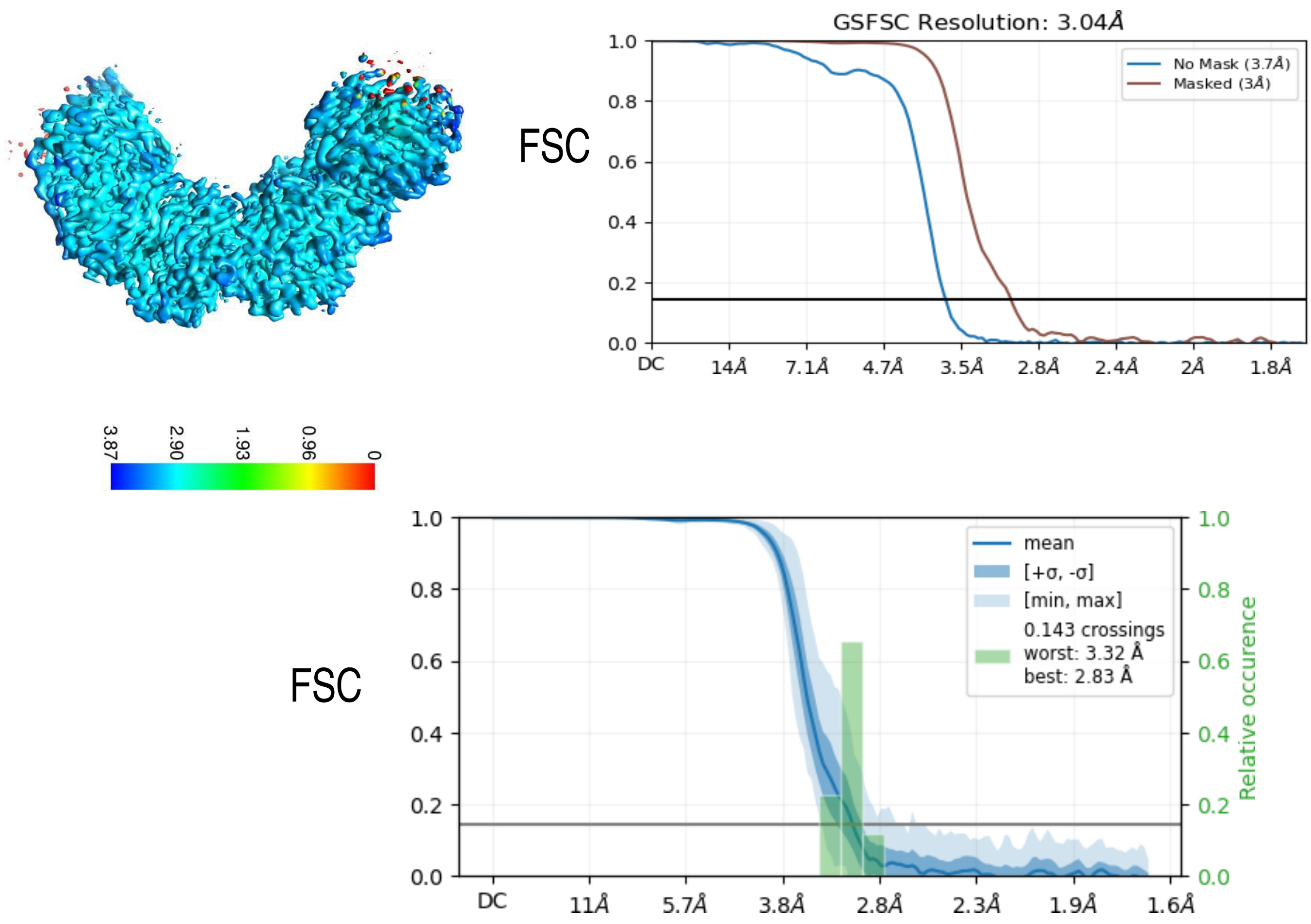

FIGURE S9: Summary of reconstruction of Cryo-EM data: a) Local resolution distribution of the reconstructed map calculated in cryosparc and colour coded. b) Gold standard Fourier Shell correlation curve of masked and unmasked map. The resolution estimates at a cutoff of 0.143 crossing indicated by solid line. c) Gold standard Fourier Shell correlation curve and the estimated mean and standard deviation. At 0.143 crossing relative occurrence of worst and best resolution distribution is indicated.

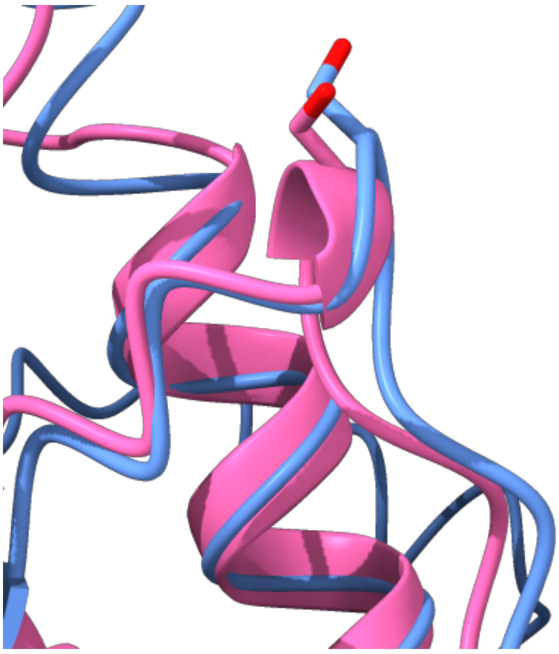

#### Figure S10: Structural alignment of *Ndas*1146 with NfsA.

*Ndas*1146 structure (pink), aligned with NfsA (blue), a homolog from the nitroreductase family. Catalytic Ser41 of NfsA (PDB: 1F5V[3]) aligns with Ser58 of Ndas1146 leading to mutational analysis of this residue to investigate potential mechanistic properties.

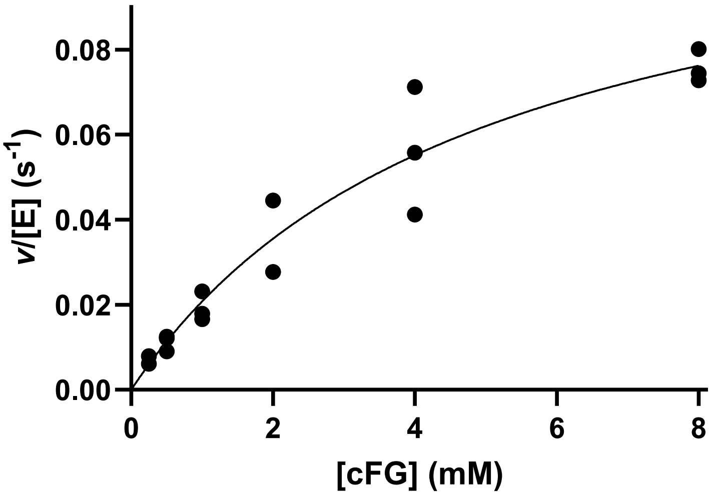

#### Figure S11: NdasCDO$_{S58A}$ kinetic analysis.

Raw data for Michaelis-Menten plots using NdasCDO-S58A mutant with cFG as a substrate.

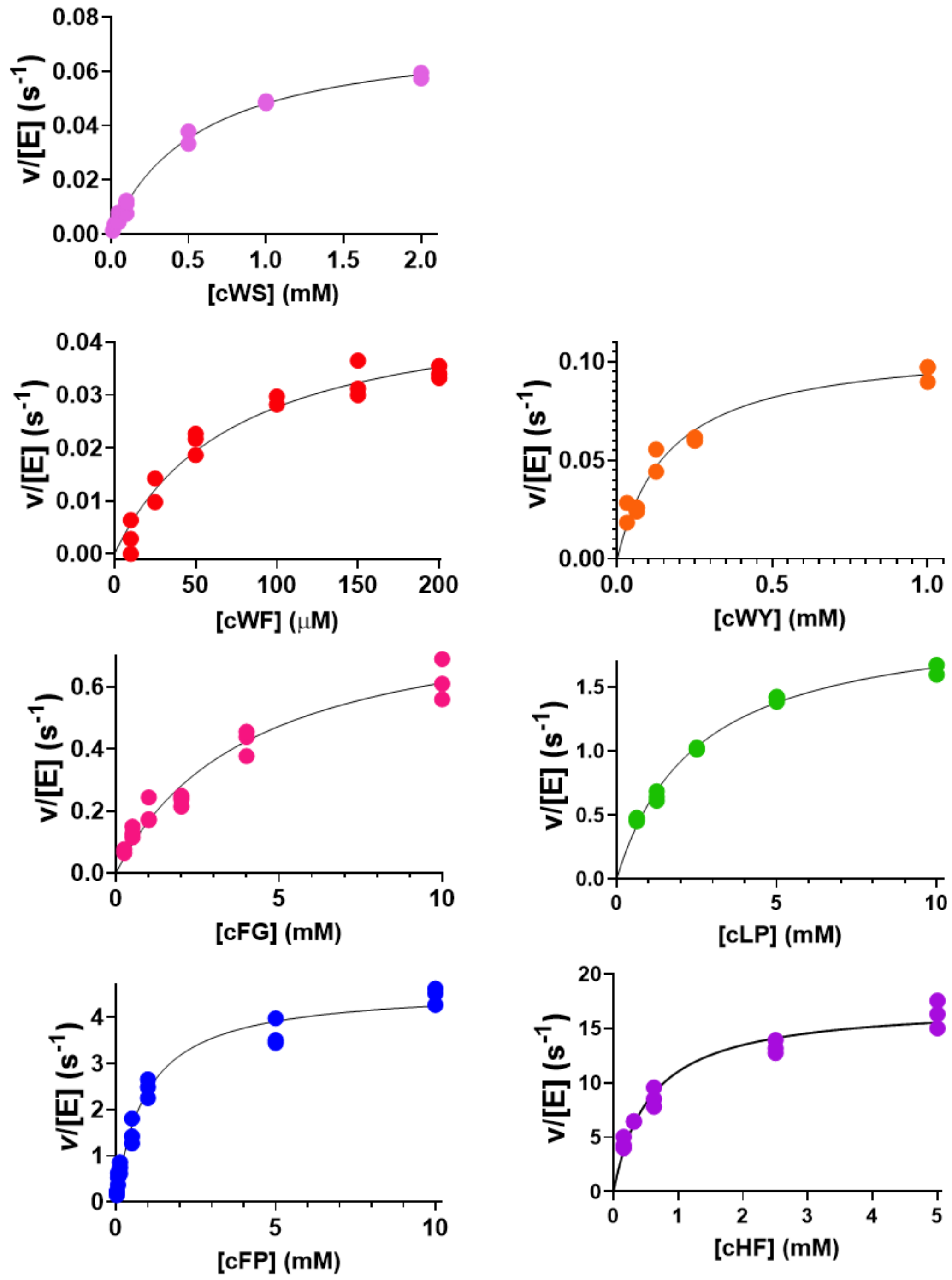

Figure S12: Raw data for data on Figure 3. Raw data for Michaelis-Menten plots with distinct substrates. All data are shown.

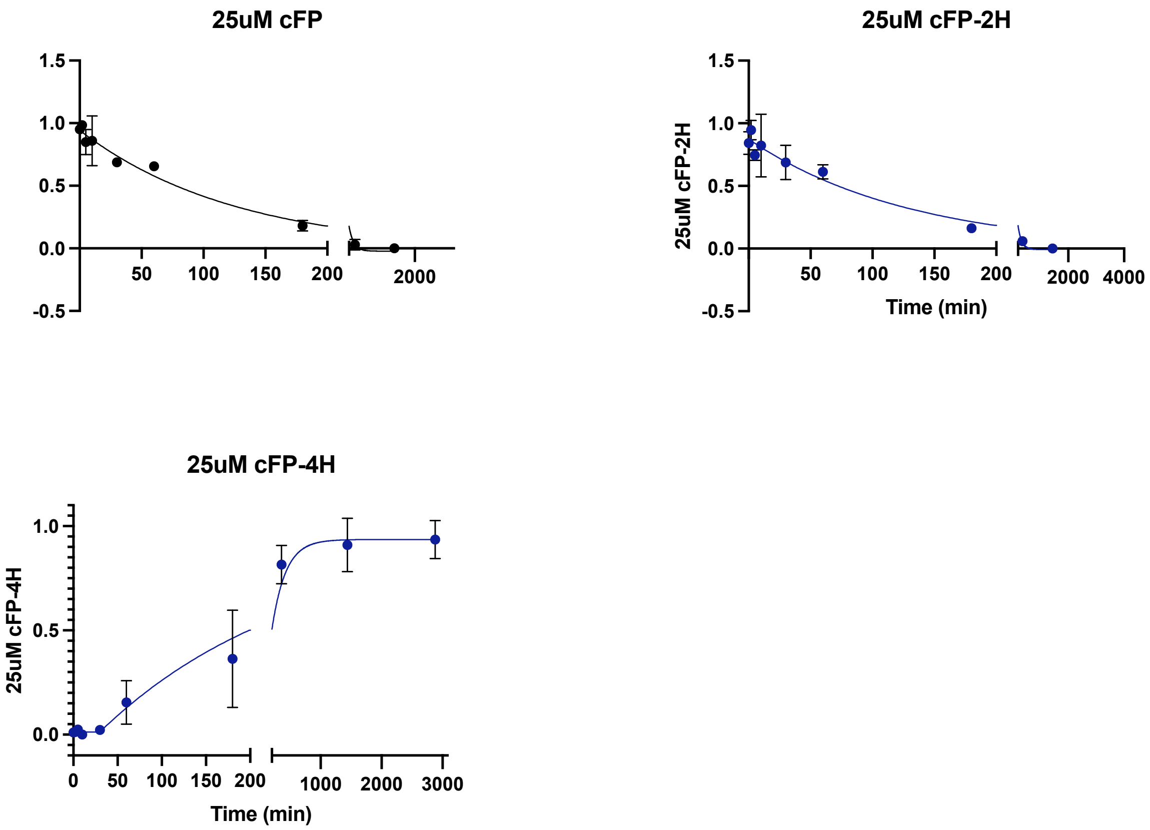

Figure S13: cFP Progress Curves. LC-MS progress curves monitoring cFP disappearance and appearance of cFP-2H and cFP-4H. Data were fitted to single exponential equations to aid data visualization. No mechanistic interpretation of rates observed is made, apart from the conclusion that there is a lag before formation of cFP-4H.

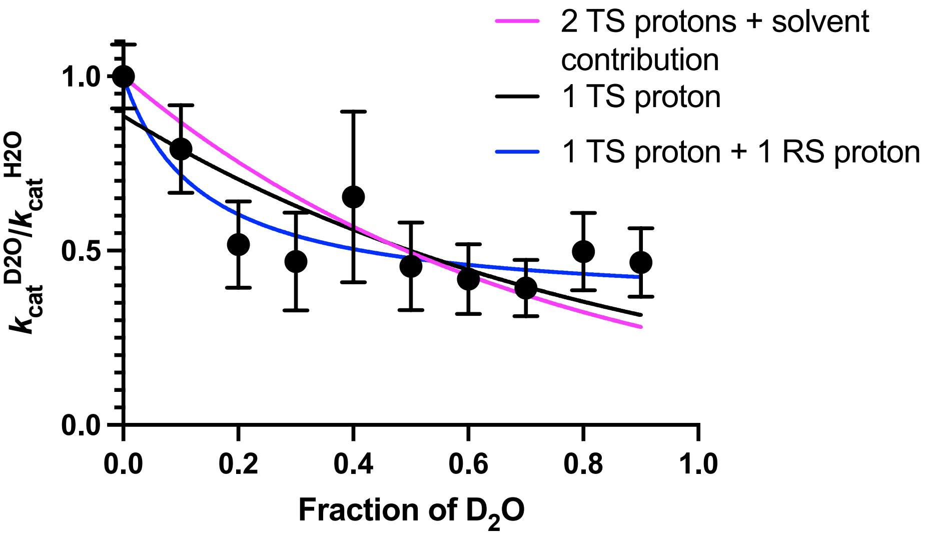

Equations for fitting:
Y= (1-x+x*ϕ1)/(1-x+x*ϕr) – 1 TS, 1 RS (blue)
Y= (1-x+x*ϕ1)^2*(ΦSolvent)^x – 2TS + bulk solvent contribution (pink)
Y= (1- x + x*ϕ1) – 1TS (black)

#### Figure S14: Proton inventories for the NdasCDO-catalyzed reaction.

Lines are fits to different models, as specified in the Figure legend. The best fitted model accounts for 1 transition state proton and 1 reactant state proton contributing to the observed $^{D2O}V = 2.1 \pm 0.1$.

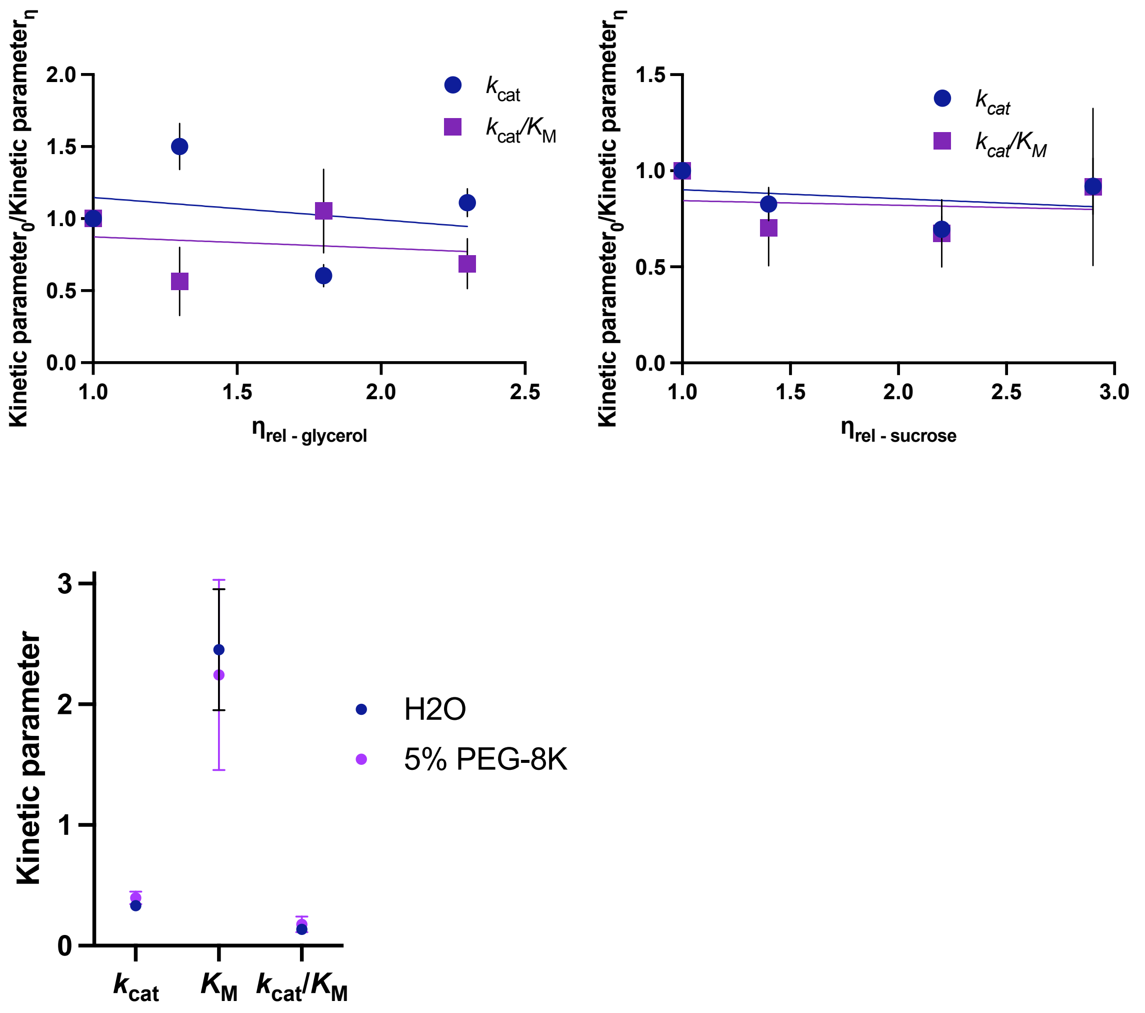

Figure S15: *Ndas*CDO Viscosity studies with glycerol, sucrose and Peg 8K.

### Supporting Tables

#### Table S1: Primers for site directed mutagenesis of *Ndas*CDO

| **Primer** | **Sequence (5'-3')** |
|---|---|
| 2-17 truncation Fwd | TCATCACACGCCCTACAG |
| 2-17 truncation Rev | CATGGTATATCTCCTTATTAAAGTTAAAC |
| 2-39 truncation Fwd | GACGACTCCCTCCTCGAC |
| 2-39 truncation Rev | CATGGTATATCTCCTTATTAAAGTTAAACAAAATTATTTC |
| S58A Fwd | CCCCTCGGCGGCCAACAAGCA |
| S58A Rev | GCGGCGAGCATGGCGTCC |

#### Table S2: Relative viscosities (ηrel) for different concentrations of glycerol and sucrose were measured by Bazelyansky *et al.* and are as below:

| **Viscogen** | **Concentration (%)** | **$\eta_{rel}$** |
|---|---|---|
| Sucrose | 14 | 1.5 |
| | 24 | 2.2 |
| | 32 | 2.9 |
| Glycerol | 10 | 1.3 |
| | 20 | 1.8 |
| | 30 | 2.3 |

#### Table S3: Data collection and refinement statistics.

| | | |
|---|---|---|
| Data collection and processing | | |
| Magnification | 105,000 | |
| Voltage(kV) | 300 | |
| Electron exposure | 33.5 | |
| Defocus Range | (1.5-3.2) | |
| Pixel size | 0.831 | |
| Symmetry imposed | C1 | |
| Initial particle images (no.) | 1,180,140 | |
| Final particle images (no.) | 849,600 | |
| Map resolution (Å) | 3.07 | |
| FSC threshold | 0.143 | |
| Map resolution range (Å) | 2.8-3.4 | |
| Helical reconstruction | 45.45 | |
| Helica Rise | 132.327 | |
| Helical Twist | 123.6 | |
| Pitch | | |
| Models | NdasA | NdasB |
| Refinement | | |
| Initial Model | Alphafold | Alphafold |
| Map sharpening B factor ($Å^2$) | 180 | 180 |
| Model Composition | | |
| Non-hydrogen atoms | 2684 | 1510 |
| Protein residues | 343 | 198 |
| Ligands | Cys-FMN | Nan |
| r.m.s deviations | | |
| Bond lengths (Å) | 0.003 | 0.08 |
| Bond Angles (°) | 0.579 | 0.825 |
| Validation | | |
| MolProbity score | 1.72 | 1.59 |
| Clashscore | 8.5 | 6.34 |
| CaBLAM outliers (%) | 2.4 | 1.05 |
| Rotamer outliers (%) | 0 | 0 |
| Cβ outliers (%) | 0 | 0 |
| Ramachandran plot | | |
| Favoured (%) | 96.12 | 96.39 |
| Allowed (%) | 3.88 | 3.61 |
| Disallowed (%) | 0 | 0 |

### Appendix 1: synthesis and characterization of cWS

#### Methodology

**Thin-layer chromatography (TLC)** analysis was conducted on pre-coated silica gel-coated 60 ($F_{254}$) and visualised with $UV_{254}$ fluorescent indicator followed by permanganate staining solution and subsequent heating for visual enhancement.

**Flash column Chromatography** utilized Merck silica gel 60 (40—63 μm) range under a positive pressure from compressed air.

**Nuclear magnetic resonance (NMR)** spectra were obtained using two distinct setups: a Bruker Advance III 500 system featuring a Prodigy BBO cryoprobe ($^{1}$H, 500 MHz; $^{13}$C, 125 MHz) and a Bruker Advance 400 system equipped with a BBFO probe ($^{1}$H, 400 MHz; $^{13}$C, 125 MHz). Spectra are referenced using the stated deuterated solvent, $^{1}$H NMR; DMSO-$d_6$ = 2.50 ppm or $CDCl_3$ = 7.26 ppm, $^{13}$C NMR; DMSO-$d_6$ = 39.5 (± 0.06) ppm or $CDCl_3$ = 77.2 (± 0.06) ppm. The reported coupling constants ($J$) were provided in Hertz, accurate to the nearest decimal place and are uncorrected. Chemical Shifts (δ) are reported in ppm (parts per million). Solvent peaks were referenced to literature values. [4]

**Optical Specific Rotations** were recorded with a Perkin Elmer Model 341 polarimeter at a wavelength of 589 nm (Sodium D line). The experiment utilized a cell with a 1 dm path length. Concentration ($c$) is expressed in g/100 mL using spectrophotometric grade methanol. Specific rotations determined at 20 °C.

**Liquid Chromatography—Mass Spectrometry (LC-MS)** (Need to collect information for that)

#### Synthetic pathway to c(WS)

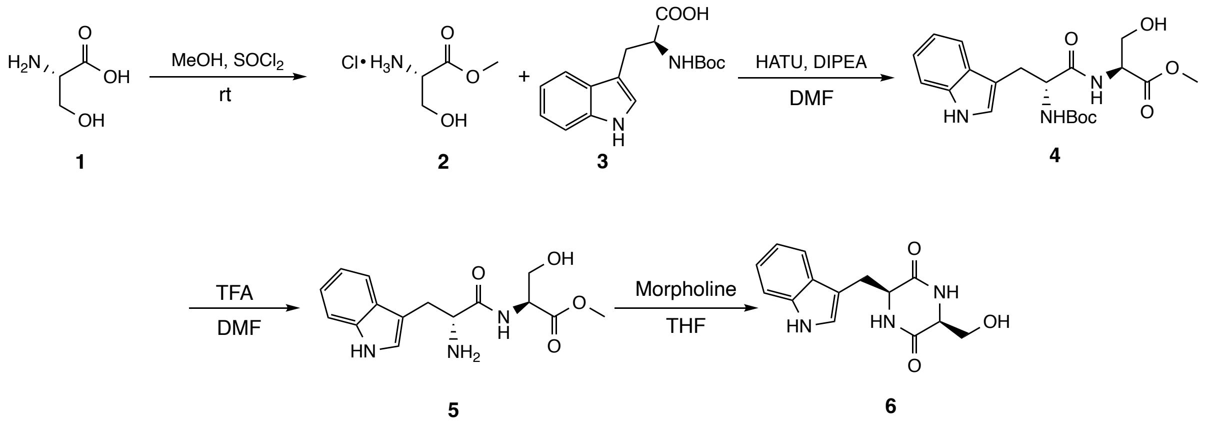

#### Methyl L-serinate (2)

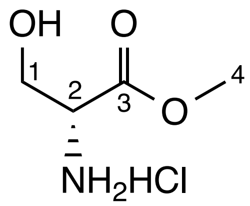

L-serine (5.00 g, 47.6 mmol) was dissolved in MeOH (64 mL) at -5 °C. Thionyl chloride (20.7 mL, 286 mmol) was slowly added over 30 minutes. After 60 hours, the mixture was concentrated, then co-evaporated with diethyl ether four times. The resulting solid was recrystallized in MeOH to give compound **2 (4.34 g, 58%)** as white crystals. Consistent with literature data. [4]

$^{1}$**H NMR** (400 MHz, DMSO-$d_6$) δ 8.64 (3H, s, $NH_3$), 5.64 (1H, s, OH), 4.08 (1H, t, $J$ = 3.5 Hz, H-2), 3.84-3.83 (2H, m, H-1), 3.74 (3H, s, H-4). $^{13}$**C NMR** (126 MHz, DMSO-$d_6$) δ 169.0, 59.9, 54.8, 53.2.

**Boc-(L-Trp-L-Ser)-OMe (22)**

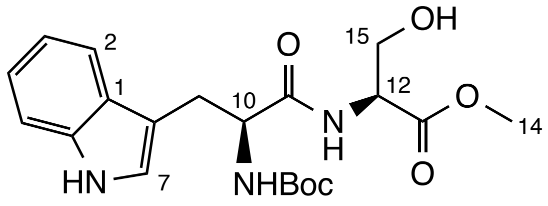

To a flame-dried flask and under nitrogen environment, Boc-tryptophan (3.00 g, 9.86 mmol), **2** (1.39 g, 8.96 mmol), HATU (3.75 g, 9.86 mmol), anhydrous DIPEA (4.7 mL, 26.9 mmol) and anhydrous DMF (20.0 mL) were added. The reaction was allowed to stir at rt until complete consumption of starting material, 16 h. The solution was extracted with ethyl acetate (3 × 30 mL). The combined extracts were washed with water (30 mL) and brine (30 mL), then dried over $MgSO_4$. Solution was then concentrated under reduced pressure, followed by co-evaporation using DCM and hexane. Purification using silica gel column chromatography (30-20% Hexanes/EtOAc) to afforded compound **4 (3.39 g, 93% yield)** as a white crystalline powder. Consistent with literature data.[5]

$R_f$ 0.35 (20% Hexanes/EtOAc); $^{1}$**H NMR** (500 MHz, $CDCl_3$) δ 8.19 (1H, s, NH), 7.67 (1H, d, $J$ = 7.9 Hz, H-2), 7.39 (1H, d, $J$ = 8.1 Hz, H-5 ), 7.23 (1H, t, $J$ = 7.5 Hz, H-4), 7.19 – 7.12 (2H, m, H-3, H-7), 6.70 (1H, d, $J$ = 6.9 Hz, NH), 5.16 (1H, s, NH ), 4.52 (1H, dt, $J$ = 7.1, 3.4 Hz, C-12), 4.43 (1H, q, $J$ = 6.7 Hz, C-10), 3.91-3.80 (2H, m, C-15), 3.72 (3H, s, C-14), 3.40 (1H, dd, $J$ = 14.6, 5.9 Hz, C-9 ), 3.22 (1H, dd, $J$ = 14.6, 6.7

Hz, C-9), 1.45 (9H, s, Boc); **$^{13}$C NMR** (126 MHz, $CDCl_3$) δ 170.3, 136.3, 127.4, 123.2, 122.5, 119.8, 118.8, 111.3, 110.4, 62.9, 55.4, 55.1, 52.7, 28.3, 27.8.

**Methyl *D*-tryptophyl-*L*-serinate (23)**

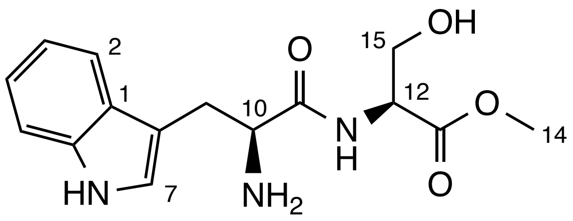

Compound **4** (3.29 g, 8.11 mmol) was added in DCM (7.0 mL) and stirred at rt until fully dissolved. TFA (3.1 mL, 40.55 mmol) was slowly added. The reaction was stirred for 16 hrs and the resulting brown solution was concentrated under reduced pressure to afford compound **5** (4.98 g) a brown viscous oil. The resulting compound was used without further purification.

***R_f*** 0.21 (20% Hexane/Ethyl acetate); **$^1$H NMR** (500 MHz, DMSO-$d_6$) δ 11.05 (1H, d, *J* = 2.5 Hz, NH), 9.09 (1H, d, *J* = 7.7 Hz, NH), 8.10 – 8.06 (2H, m, $NH_2$), 7.76 (1H, d, *J* = 7.9 Hz, H-2), 7.38 (1H, d, *J* = 8.2 Hz, H-5), 7.24 (1H, d, *J* = 2.4 Hz, H-7), 7.11 (1H, t, *J* = 7.5 Hz, H-4), 7.02 (1H, t, *J* = 7.5 Hz, H-3), 4.47 (1H, dt, *J* = 7.6, 4.6 Hz, H-12), 4.14 (1H, dt, *J* = 9.5, 5.3 Hz, H-10), 3.81 (1H, dd, *J* = 11.1, 4.9 Hz, H-15), 3.69 – 3.66 (4H, m, H-15/H-14), 3.29 (1H, dd, *J* = 14.7, 5.4 Hz, H-9), 3.07 (1H, dd, *J* = 14.8, 8.7 Hz, H-9).

**Cyclic(L-trp-L-ser) (24)**

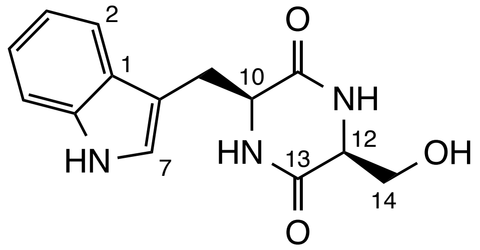

Compound **5** (2.28 g, 7.47 mmol) was dissolved in THF (21.0 ml) and stirred vigorously. The solution was brough to 0 °C. Subsequently, morpholine (9.1 ml, 104.54 mmol) was added dropwise. The solution was allowed to warm up to rt and left to stir for 48 hr. It was then concentrated under reduced pressure and purified using flash column chromatography (1-20% MeOH/DCM) to offer compound **6** (0.53 g, 26%, over 2 steps) as a white powder. Consistent with literature data.[6]

***R_f*** 0.19 (10% methanol/DCM); $[\alpha]_D^{20}$ –99.6 (*c* 0.36, MeOH); **$^1$H NMR** (500 MHz, DMSO-$d_6$) δ 10.88 (1H, s, NH-Indole), 7.89 (2H, m, NH x 2), 7.54 (1H, d, *J* = 7.9 Hz, H-2), 7.34 (1H, d, *J* = 8.1 Hz, H-5), 7.13 (1H, d, *J* = 2.4 Hz, H-7), 7.06 (1H, t, *J* = 7.5 Hz, H-4), 6.96 (1H, t, *J* = 7.4 Hz, H-3), 4.91 (1H, t, *J* = 5.6 Hz, OH), 4.02 (1H, dt, *J* = 6.9, 3.6 Hz, H-10), 3.68 (1H, dt, *J* = 5.7, 2.8 Hz, H-12), 3.33 – 3.28 (1H, m, H-$14^a$), 3.19 (2H, qd, *J* = 14.4, 5.6 Hz, H-9), 3.05 (1H, dt, *J* = 10.9, 5.6 Hz, H-$14^b$). **$^{13}$C NMR** (126 MHz, DMSO-$d_6$) δ 167.7, 166.2, 136.5, 128.1, 124.6, 121.3, 119.1, 118.8, 111.7, 109.6, 63.5, 57.8, 56.0, 30.9.

### Selected $^{1}H$ and $^{13}C$ NMR spectra

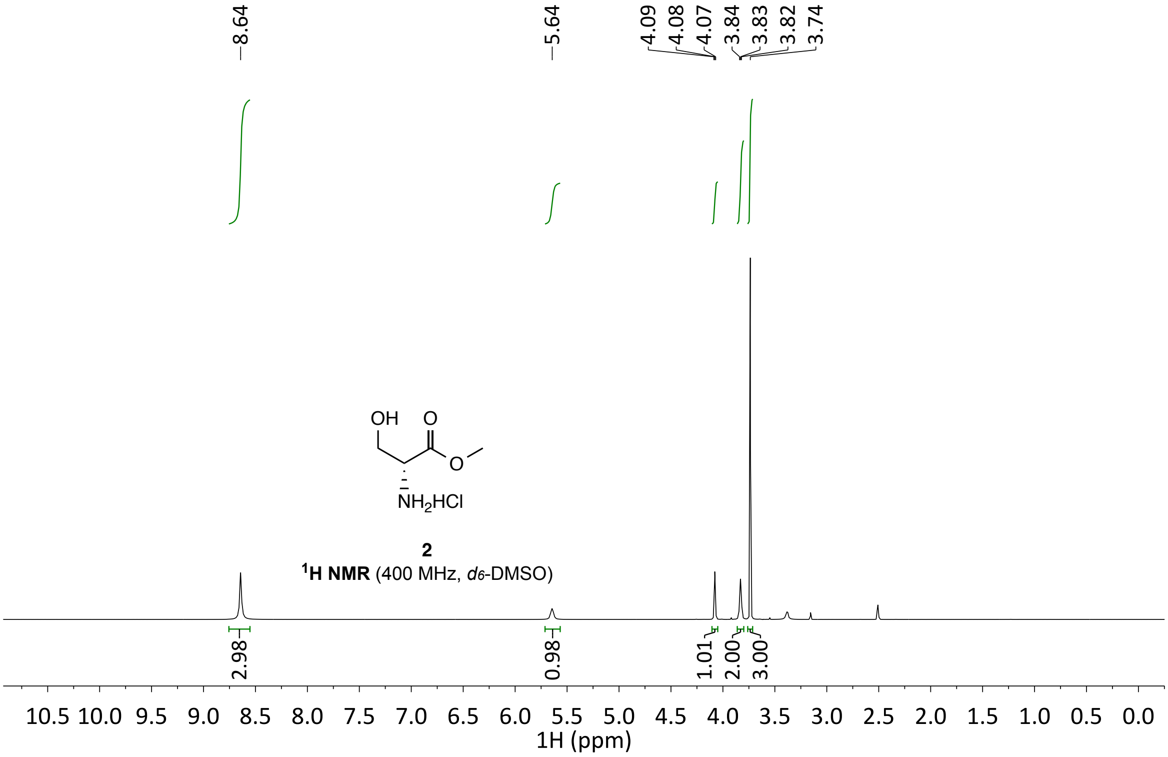

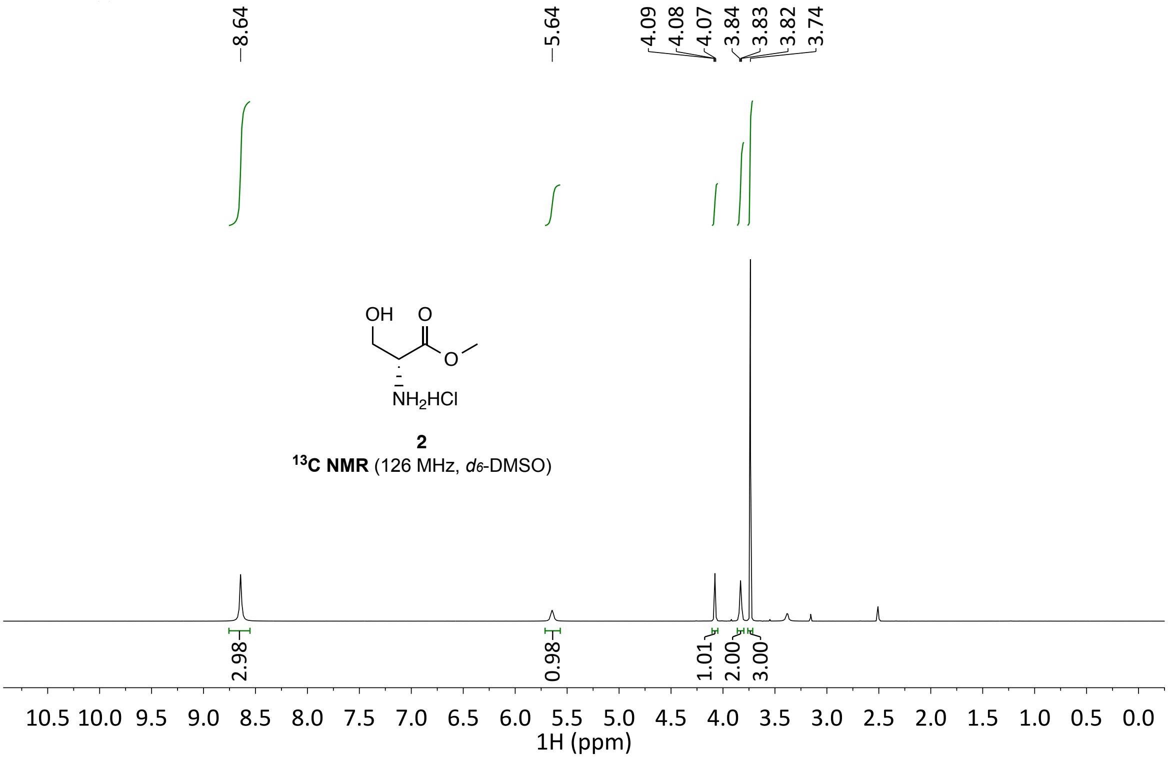

**2**
**$^{13}$C NMR** (126 MHz, *d6*-DMSO)

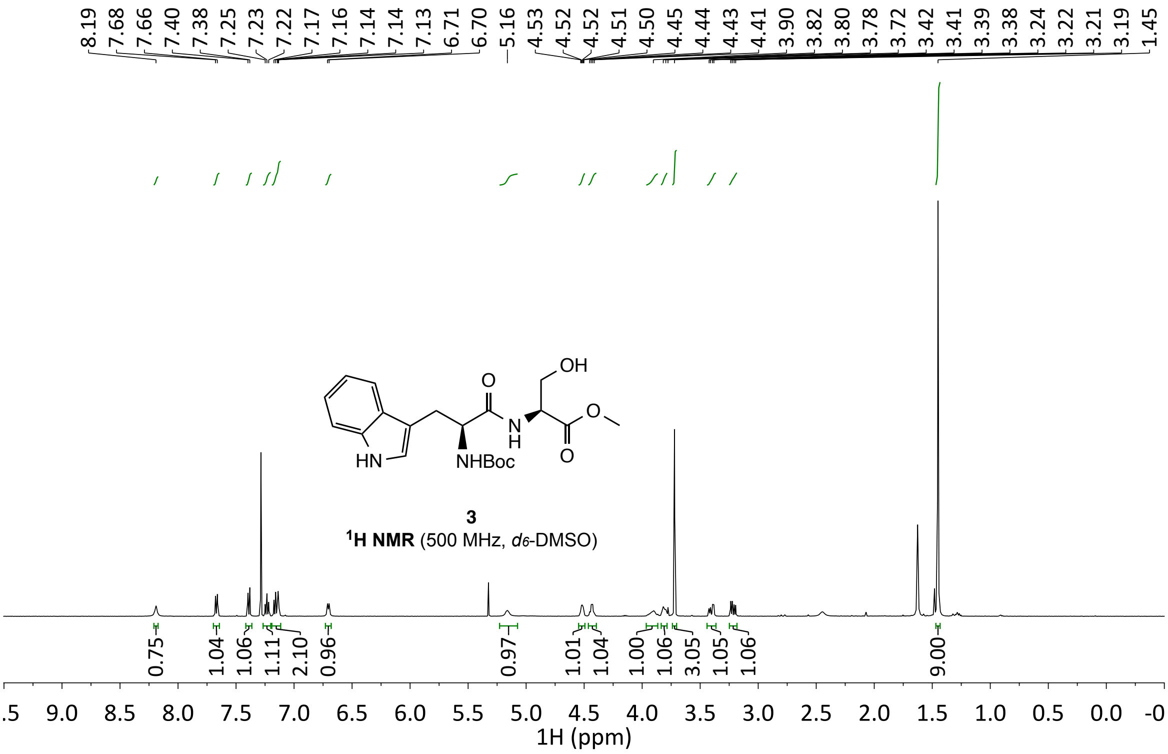

**3**
**$^{1}$H NMR** (500 MHz, *d6*-DMSO)

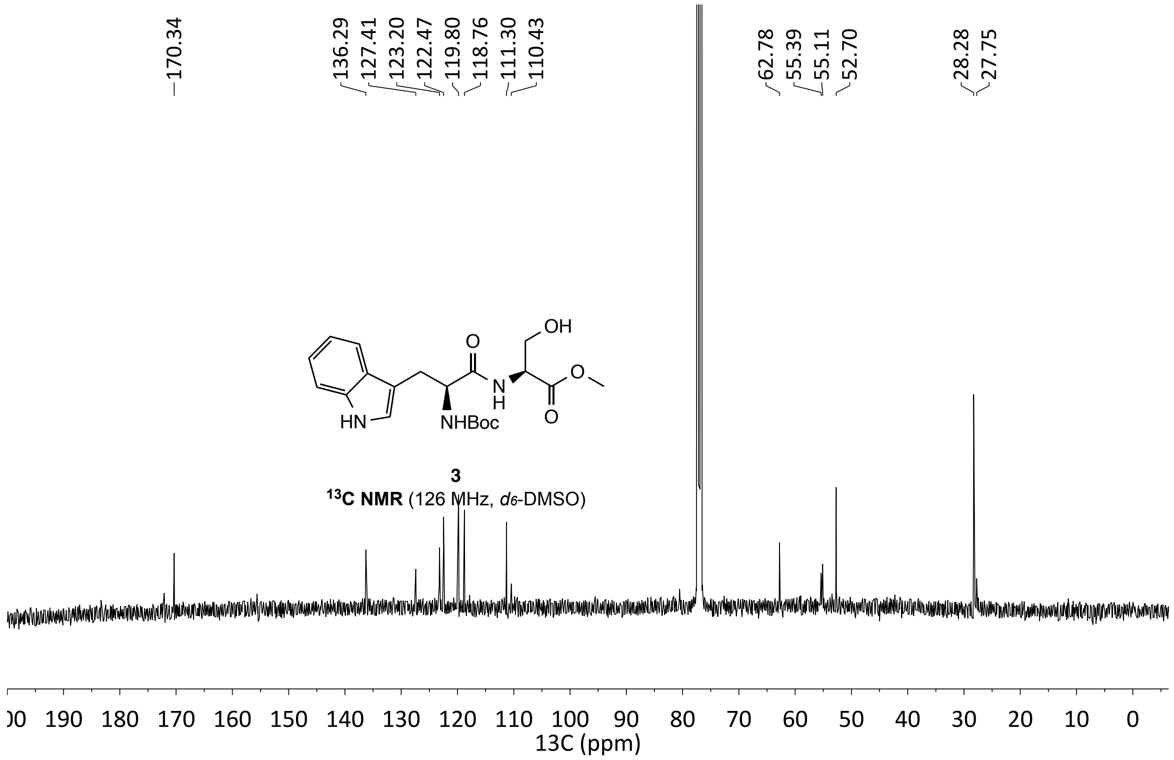
170.34
136.29
127.41
123.20
122.47
119.80
118.76
111.30
110.43
62.78
55.39
55.11
52.70
28.28
27.75
OH
O
N
H
O
O
HN
NHBoc
3
13C NMR (126 MHz, d6-DMSO)
00 190 180 170 160 150 140 130 120 110 100 90 80 70 60 50 40 30 20 10 0
13C (ppm)

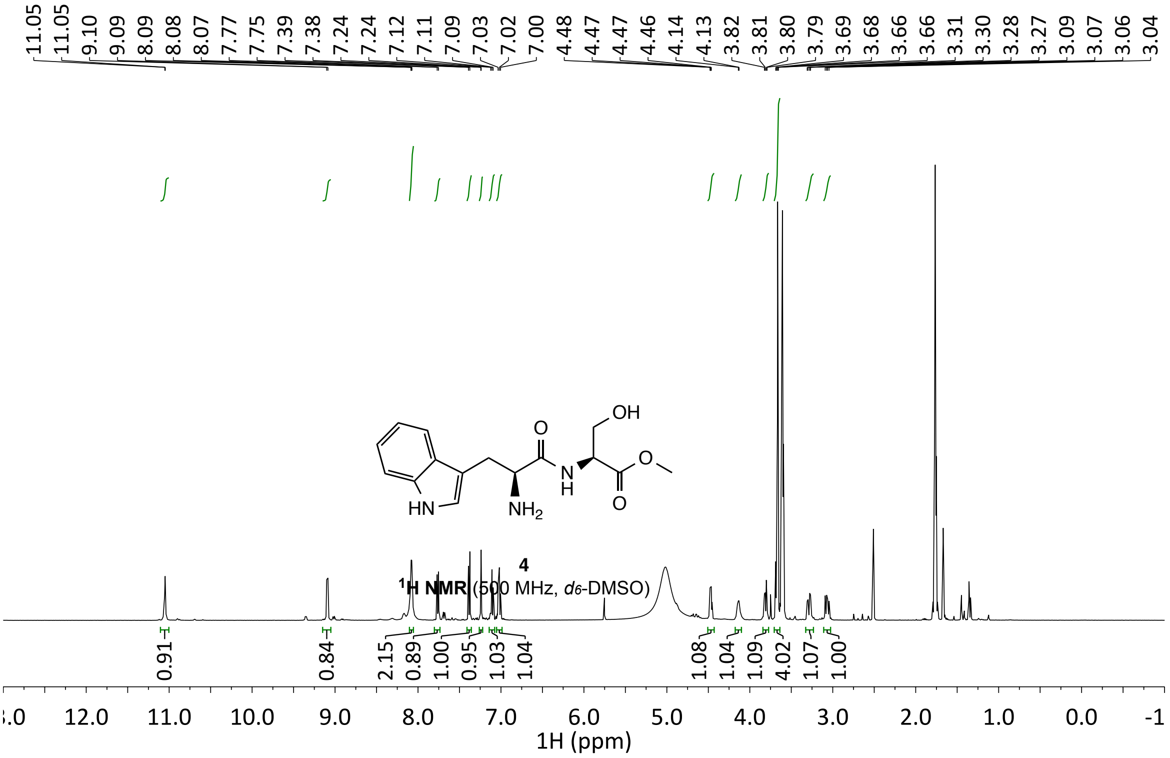
11.05 11.05 9.10 9.09 8.09 8.08 8.07 7.77 7.75 7.39 7.38 7.24 7.24 7.12 7.11 7.09 7.03 7.02 7.00 4.48 4.47 4.47 4.46 4.14 4.13 3.82 3.81 3.80 3.79 3.69 3.68 3.66 3.66 3.31 3.30 3.28 3.27 3.09 3.07 3.06 3.04
OH
O
N
H
O
O
HN
NH2
4
1H NMR (500 MHz, d6-DMSO)
0.91 0.84 2.15 0.89 1.00 0.95 1.03 1.04 1.08 1.04 1.09 4.02 1.07 1.00
12.0 11.0 10.0 9.0 8.0 7.0 6.0 5.0 4.0 3.0 2.0 1.0 0.0
1H (ppm)

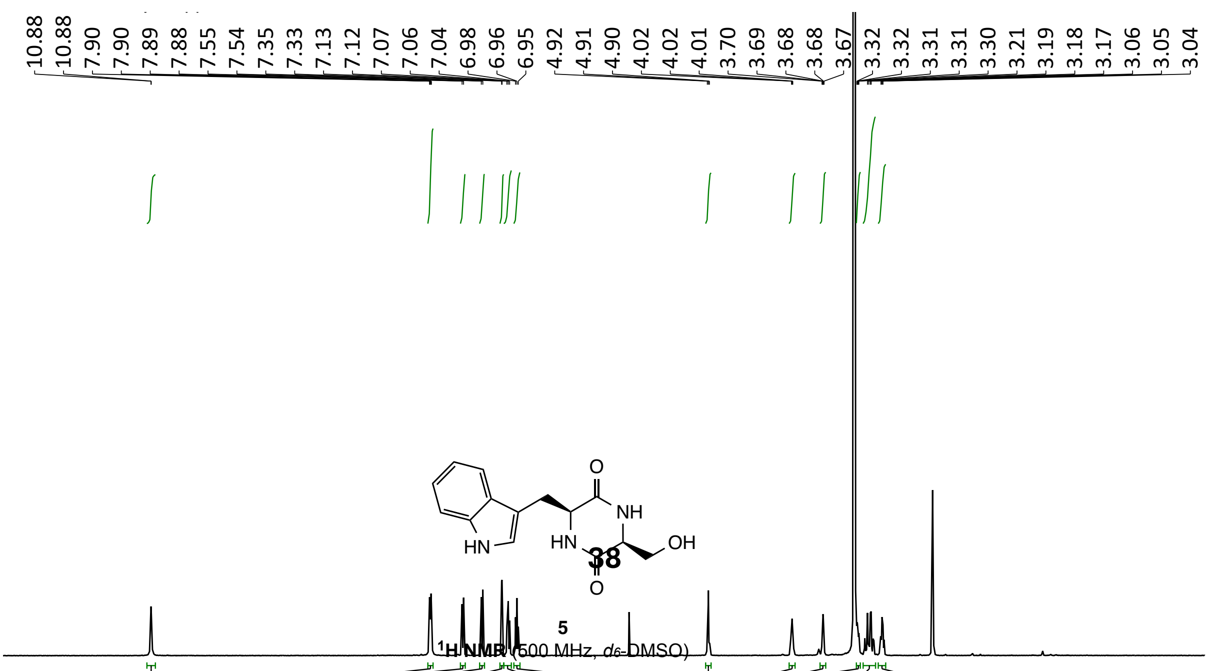
10.88 10.88 7.90 7.90 7.89 7.88 7.55 7.54 7.35 7.33 7.13 7.12 7.07 7.06 7.04 6.98 6.96 6.95 4.92 4.91 4.90 4.02 4.02 4.01 3.70 3.69 3.68 3.68 3.67 3.32 3.32 3.31 3.31 3.30 3.21 3.19 3.18 3.17 3.06 3.05 3.04
O
NH
HN
OH
O
HN
5
1H NMR (500 MHz, d6-DMSO)

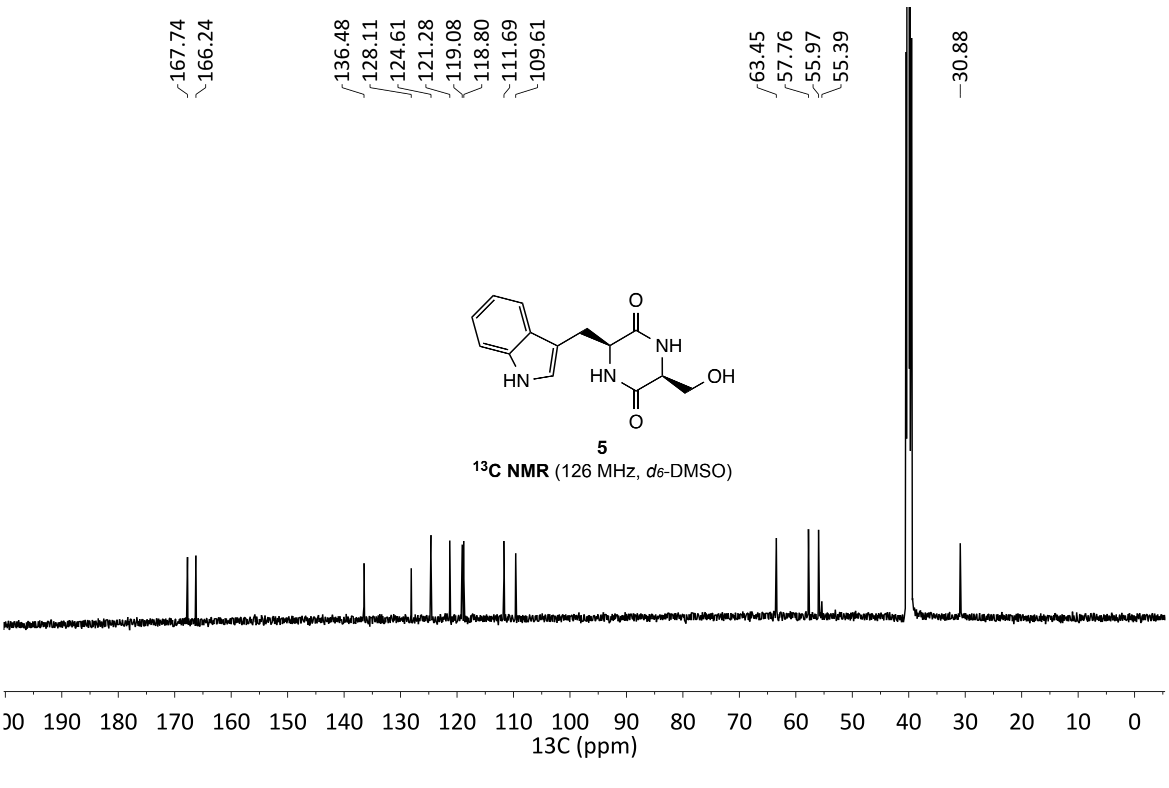
167.74
166.24
136.48
128.11
124.61
121.28
119.08
118.80
111.69
109.61
63.45
57.76
55.97
55.39
30.88
O
NH
HN
HN
OH
O
5
$^{13}$C NMR (126 MHz, $d_6$-DMSO)
190 180 170 160 150 140 130 120 110 100 90 80 70 60 50 40 30 20 10 0
13C (ppm)

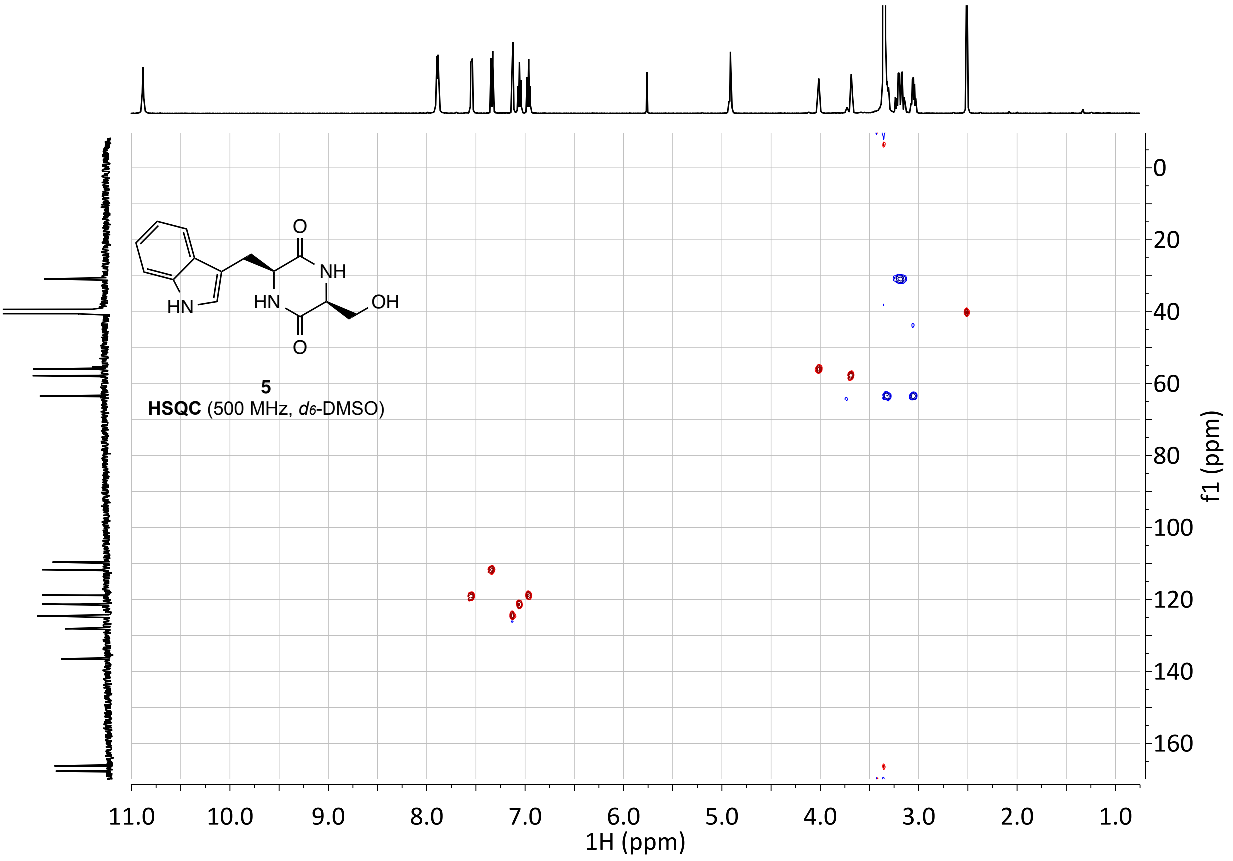
HN
HN
NH
O
O
OH
5
HSQC (500 MHz, d6-DMSO)
1H (ppm)
f1 (ppm)

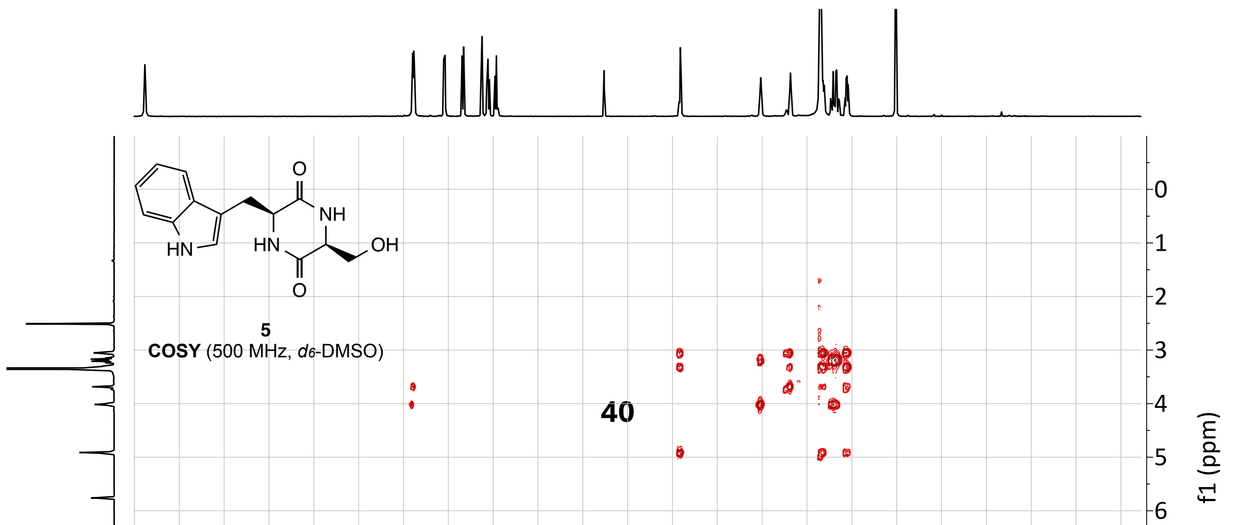
HN
HN
NH
O
O
OH
5
COSY (500 MHz, d6-DMSO)
f1 (ppm)

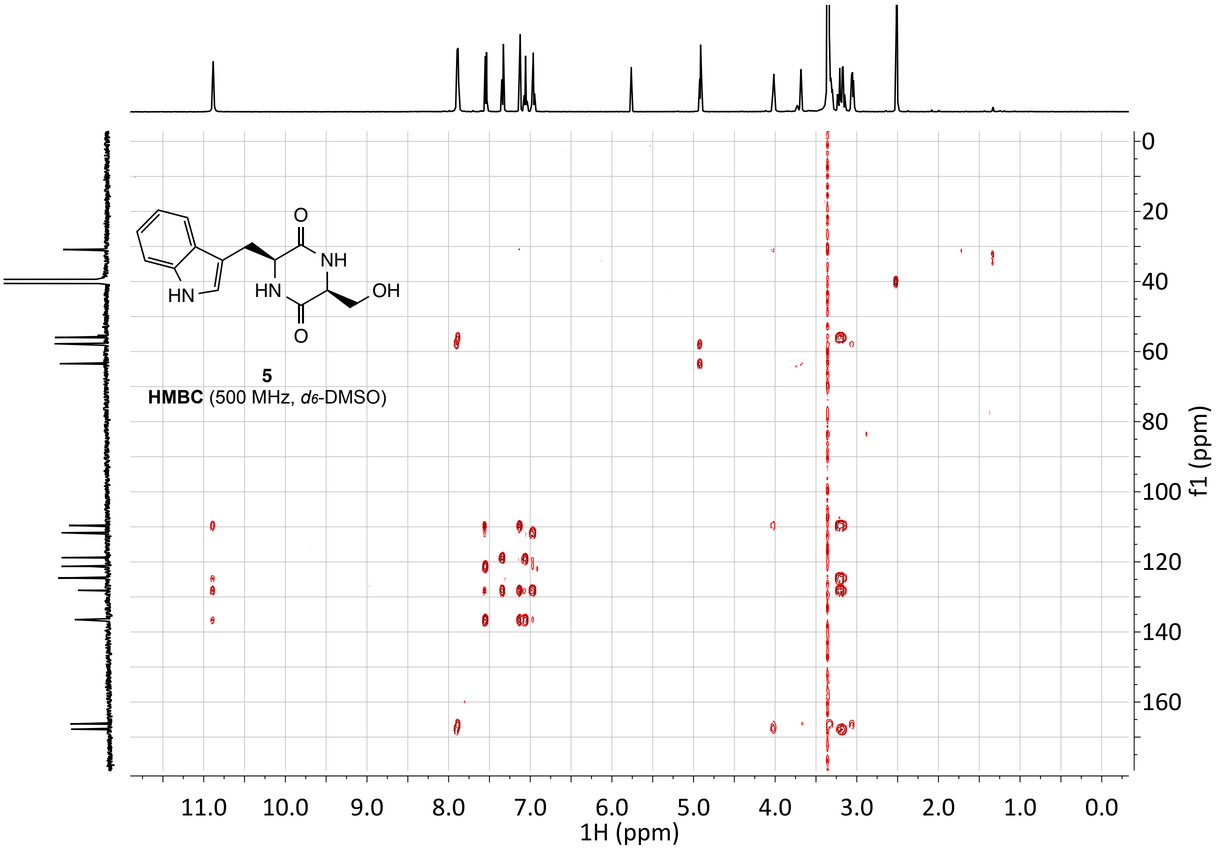
O
NH
HN
OH
HN
O
5
HMBC (500 MHz, d6-DMSO)
11.0
10.0
9.0
8.0
7.0
6.0
5.0
4.0
3.0
2.0
1.0
0.0
1H (ppm)
0
20
40
60
80
100
120
140
160
f1 (ppm)
